# Interindividual differences in caries potential of the salivary microbiome after mouthwash exposure

**DOI:** 10.1101/2025.01.13.632702

**Authors:** Fabian Mermans, Britt Poelman, Mehraveh Saghi, Wim Teughels, Nico Boon

**Author notes:** Correspondence to: Nico Boon, Ghent University, Faculty of Bioscience Engineering; Center of Microbial Ecology and Technology (CMET); Frieda Saeysstraat 1, B-9052 Gent, Belgium.

## Abstract

Prevention and treatment of dental caries is often achieved by fluoride mouthwashes. The most common forms of fluoride are Stannous fluoride (SnF_2_), amine fluoride (AmF) and sodium fluoride (NaF), each with differing activities against oral bacteria. Additionally, differences in oral microbiome between individuals are linked to differences in response to treatment and the phenotype of the microbiome can provide information on this response. Flow cytometry can successfully track microbial phenotypic heterogeneity and may prove useful for capturing the response to treatment. In this study, we compared the effect of three fluoride-containing mouthwashes, Elmex Anti Caries (AmF/NaF), Listerine Anticaries (NaF) and Meridol (AmF/SnF_2_), on the salivary microbiome and assessed their anticaries potential. Furthermore, we investigated if the flow cytometric fingerprint could be used to capture the response to treatment. Saliva samples of different donors were treated with the mouthwashes and the intact salivary community composition was assessed using 16S rRNA gene amplicon sequencing. Treated saliva samples were incubated in sucrose-rich medium to assess the organic acid production of the remaining microbiota. Moreover, flow cytometric fingerprints of the intact microbial population were used to track the response to treatment and to train a random forest classifier predicting the most suitable mouthwash for the donor. We found a mouthwash-dependent shift in the salivary microbiome composition and found different anticaries potential of the mouthwashes. Meridol affected more bacterial taxa and had the best overall anticaries potential, followed by Elmex. Finally, the response to treatment could be captured in the flow cytometric fingerprint of the initial intact salivary microbiota and a random forest classifier predicting the most suitable mouthwash for the donor was successfully trained.

## Introduction

The highly diverse microbial community in the oral cavity is structured in polymicrobial biofilms on the teeth, subgingival surfaces and tongue, and has a wide range of metabolic capabilities [1]. These metabolic capabilities are influenced by the microbial community composition, the availability of oxygen, and the availability of nutrients supplied from saliva, the crevicular fluid and nutrition [2]. Because of the short transit time of food in the mouth, oral bacteria use free sugars and fermentable carbohydrates as their primary source of carbon [3,4]. When there is overexposure to dietary carbohydrates, in combination with host factors, production of extracellular polymeric substances (EPS) and organic acids from carbohydrate metabolism is promoted, and accumulation of acidogenic and aciduric microorganisms occurs. The increased concentration of organic acids in the tooth-associated biofilm can lower the pH below 5.5, which leads to tooth demineralization and eventually to tooth decay [4–9]. This pathology, also known as dental caries, is estimated to affect 2.5 billion people globally [10]. In a symbiotic state at near neutral pH, the oral microbial communities are dominated by *Actinomyces*, *Bifidobacteria*, *Fusobacteria*, *Neisseria*, *Porphyromonas*, non-mutans streptococci and *Veillonella* [6,11,12]. When caries develops and the acidogenic and aciduric species are promoted, the dominance of *Veillonella*, *Neisseria*, *Streptococcus*, *Propionibacterium*, *Prevotella*, and *Lactobacillus* increases [6,11,13,14].

Fluoride mouthwashes are commonly used to prevent dental caries [15–19]. Topical administration of fluoride provides its anti-cariogenic effects by inhibiting demineralization, promoting remineralization and inhibiting plaque bacteria [20]. Inhibition of demineralization is accomplished through adsorption of fluoride into the surface enamel crystal structure, forming fluorapatite, which is less prone to acid dissolution than normal tooth minerals (hydroxyapatite and carbonated apatite) [20,21]. Remineralization is promoted by acceleration of fluorapatite crystal growth in carious lesions. When fluoride adsorbs into the surface enamel, it attracts calcium ions present in saliva, which in turn attract more fluoride [20]. The inhibition of plaque bacteria is achieved by cellular uptake of fluoride ions under acidic conditions. This results in inhibition of growth, metabolic activity and enzymatic activity, thereby hindering the bacteria’s ability to form plaque [22–25]. The use of fluoride compounds in oral care products has been reported to induce a shift in microbial community composition, leading to a proportional decrease of disease-associated bacteria such as members of the *Streptococcus* and *Prevotella* genera. Thus, these compounds put a selective pressure on the oral microbiota in favor of the health-associated taxa [26–32]. Different fluoride-containing ingredients are used in mouthwashes of which the most common forms are stannous fluoride (SnF_2_), amine fluoride (AmF) and sodium fluoride (NaF). Multiple reports indicate that SnF_2_ and AmF are more effective in reducing erosive enamel loss and have more favorable effects on other clinical oral health parameters such as the approximal plaque index compared to NaF [33–36]. Additionally, higher reduction of pathogenic bacteria and better inhibition of oral microbial biofilm formation have been reported for SnF_2_ and AmF compared to NaF [31,34,37,38]. These findings suggest that the antimicrobial effect of the oral care products is mostly attributable to the fluoride counter ions and additional ingredients in the products instead of the fluoride ions themselves [39]. Indeed, the *in vivo* importance of the antimicrobial activity of fluoride in caries inhibition is questioned [23,39].

Individual differences in oral microbiome composition are significant, and previous research has demonstrated that treatment responses are associated with these interindividual variations in microbiome composition [40–42]. As such, the oral microbiome serves as a valuable source of information for predicting a patient’s response to treatment. Typically, the community composition derived from next-generation sequencing (NGS) is used as a “microbial signature” for patients [43,44]. However, this genotypic fingerprint only identifies the microorganisms present and does not provide insights into the functional phenotype of the microbial community. Because the phenotype of the microbial community can also influence treatment responsiveness, it is crucial to include phenotypic data when evaluating treatment outcomes [45,46]. Flow cytometry has proven particularly relevant when assessing phenotypic heterogeneity in microbial populations and has already been used to track the salivary microbiome, making it a fitting tool to study the phenotype of oral microbial communities [47–52].

Interestingly, no studies have directly compared the effects of different fluoride-containing ingredients on the oral microbiome. To address this gap, we investigate the impact of various fluoride-containing mouthwashes on the salivary microbiome. Given the selective pressures fluoride exerts on oral bacteria, we hypothesize that these treatments will induce shifts in microbiome composition. Furthermore, we anticipate that both the changes in microbiome composition and the anticaries effects of the mouthwashes will depend on the specific fluoride formulation. Additionally, we explore whether the anticaries effects and microbial responses to treatment are donor-dependent. To this end, we examined whether both phenotypic and genotypic fingerprints of the salivary microbiome could capture treatment responses. Saliva samples from different donors were treated with three distinct fluoride-containing mouthwashes. Following treatment, we analyzed the composition of the intact microbial cell population using 16S rRNA gene amplicon sequencing and constructed phenotypic fingerprints with flow cytometry. Finally, we assessed the organic acid production of the treated microbial populations by incubating the samples in minimal medium with sucrose as the carbon source.

## Materials and Methods

A general overview of the workflow is provided in Figure 1.

**Figure 1.**
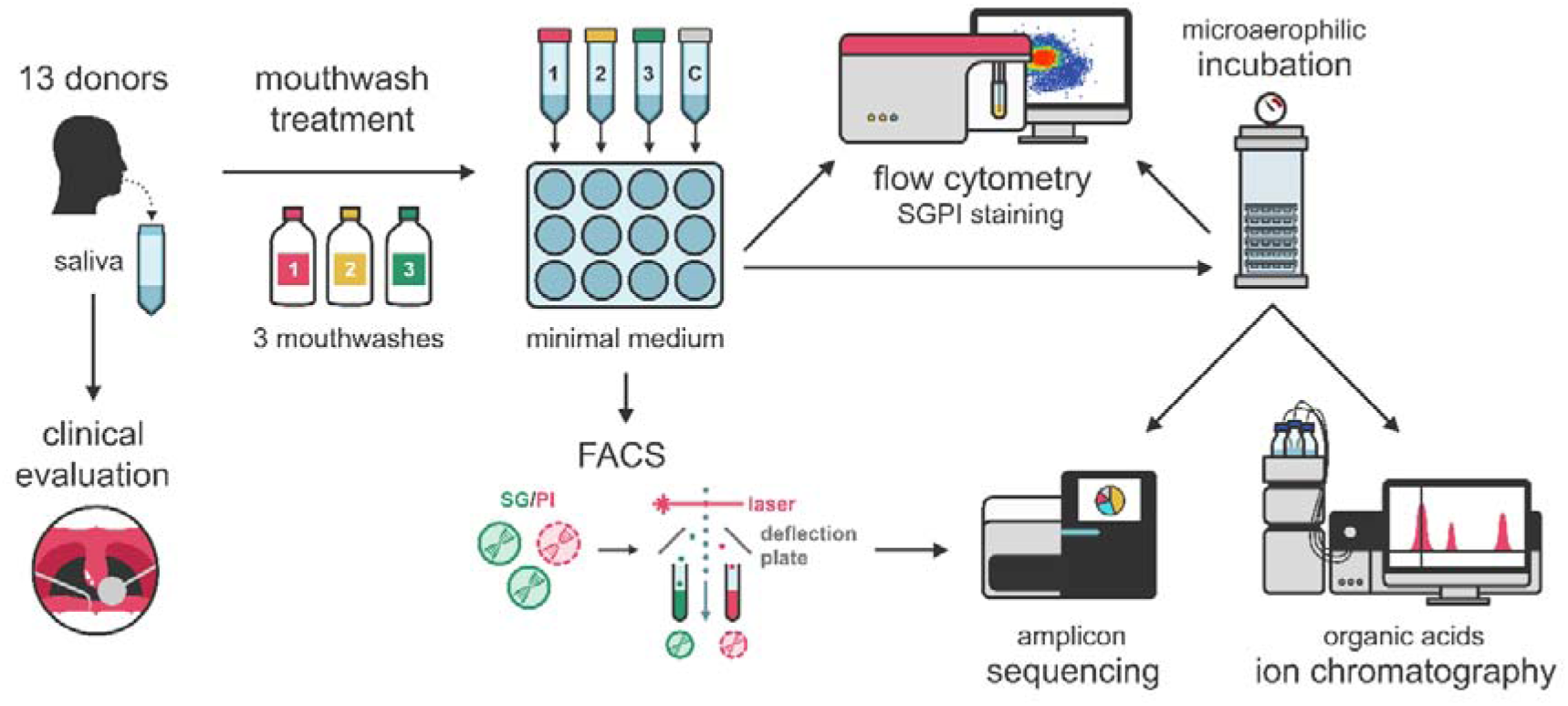
Overview of the experimental workflow. Individuals donated saliva through passive drooling. After saliva donation, donors underwent a clinical evaluation of their oral health. Saliva was used for treatment with three different kinds of mouthwashes (Elmex Anti Caries, Listerine Anti Caries and Meridol). A fourth aliquot was treated with phosphate buffered saline (PBS) and served as a control. After treatment, samples were analyzed using flow cytometry and the intact cell population was sorted (FACS) for 16S rRNA gene amplicon sequencing. Additionally, the samples were used for microaerophilic incubation in unbuffered minimal medium at 37°C. After 19h of incubation, samples were taken for 16S rRNA gene amplicon sequencing, flow cytometry and organic acids (C1-C5) were determined through ion chromatography.

### Saliva sampling and clinical evaluation of oral health status

A volume of 5 mL of saliva was collected by passive drooling from 13 individuals that did not undergo any antibiotic treatment three months prior to donation (Supplementary 1). Donors were asked to refrain from brushing their teeth, eating and drinking for two hours before donation. Moreover, donors were instructed not to use mouthwash on the day of donation. After donation, a dentist performed a clinical evaluation of the oral cavity. Teeth were examined for decay, restorations, crowns, fixed orthodontic retainers, bridges, extracted teeth and implants. Probing pocket depth (PPD) of the groove between gums and teeth was measured, as well as bleeding on probing (BOP) of the gums, gum recession and clinical attachment loss (CAL). Additionally, donors were asked to chew on plaque-disclosing tablets for 30 seconds and rinse with water afterwards to evaluate plaque level (Supplementary 1). Prior to donation and clinical evaluation, donors signed an informed consent form. The study was approved by the Ethics Committee of the University of Ghent (B6702022000406).

### Treatment and incubation of saliva

Before treatment, 500 µL of saliva was transferred to a microcentrifuge tube and samples were centrifuged at 20238 x g for 10 minutes. Supernatant was discarded and the remaining pellet was stored at -20°C up until DNA extraction. The remaining saliva was used for treatment with three different mouthwashes for 1 minute in triplicate: Listerine Anti Caries (NaF, 220 ppm F^-^) (Johnson & Johnson, Maidenhead, United Kingdom), Elmex Anti Caries (AmF and NaF, 250 ppm F^-^) (Colgate-Palmolive, Świdnica, Poland) and Meridol (AmF and SnF_2_, 250 ppm F^-^) (Colgate-Palmolive, Świdnica, Poland). A detailed list of the ingredients of the mouthwashes is listed in Supplementary 2. For donors 1-5 a mouthwash concentration of 9% v/v was used. For donor 6-13 a concentration of 7% v/v was used so that a larger proportion of the microbial community would survive the treatment (Supplementary 9; Figure S9). Moreover, control samples were included in triplicate by treating saliva with sterile phosphate buffered saline (PBS) (PBS tablet, Sigma-Aldrich, Steinheim, Germany) in the same concentration as the mouthwashes. Treatment was stopped by diluting 100-fold in unbuffered minimal medium before putting the samples on ice. The unbuffered minimal medium (Table 1) was prepared by adding 20 mL/L vitamin mix and 20 mL/L sucrose-vitamin C mix to the base medium (pH7) [11].

**Table 1.**
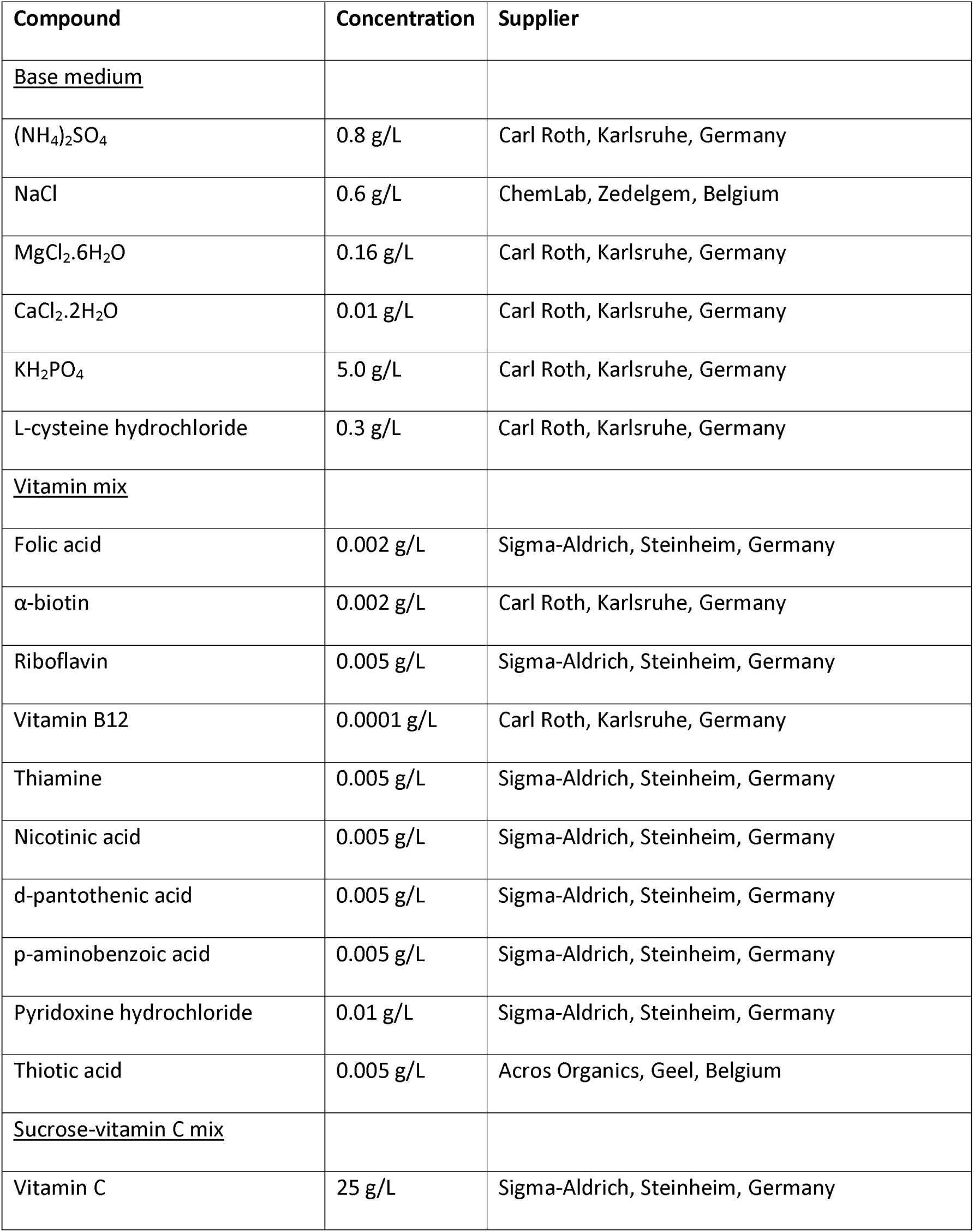
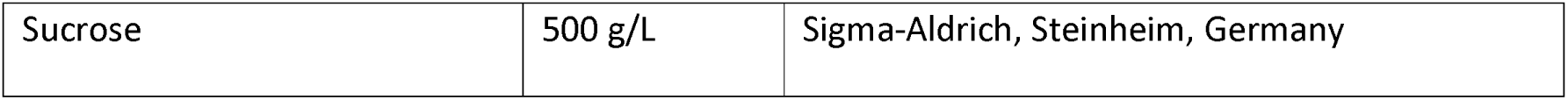
Composition of the individual components of unbuffered minimal medium for saliva as described by McLean et al. [11]. The unbuffered minimal medium was prepared by adding 20 mL/L of both the vitamin mix and the sucrose-vitamin C mix to the base medium. The pH of the base medium was adjusted to 7 before adding the respective mixes.

After treatment, the pH of the samples was determined (Supplementary 3) and samples were taken for ion chromatography (IC), flow cytometry and FACS. Additionally, 3 mL of each replicate was transferred to a sterile 12-well plate with a lid for each donor. 3 mL of unbuffered minimal medium was added to a 12-well plate as an abiotic control as well. Consecutively, 12-well plates were incubated under microaerophilic conditions (5-10 % O_2_) at 37°C for 19 hours. After incubation, the pH was determined (Supplementary 3) and samples were taken for ion chromatography and flow cytometry. Furthermore, 500 µL of sample was centrifuged at 20238 x g for 10 minutes; supernatant was discarded, and the remaining pellet was stored at -20°C for DNA extraction.

### Flow cytometry and FACS

#### Sample preparation

After treatment, an aliquot of each sample was diluted 10-fold in sterile PBS (PBS tablet, Sigma-Aldrich, Steinheim, Germany). Next, samples were stained with SYBR® Green I/propidium iodide (SGPI) to assess membrane integrity [54]. A SGPI stock solution was prepared by diluting SYBR® Green I (10 000x concentrate in DMSO) (Invitrogen, Eugene, USA) and propidium iodide (20mM) (Invitrogen, Eugene, USA) 100 and 50 times respectively, in 0.2 µm filtered DMSO (Merck, Darmstadt, Germany). Samples were then stained with 1% v/v SGPI stock solution and incubated in the dark at 37°C for 20 minutes [55].

#### Sample measurement

Stained samples were measured using an Attune NxT (Invitrogen, Carlsbad, USA) flow cytometer equipped with a blue (488 nm) and red (638 nm) laser. Performance of the instrument was checked using Attune Performance tracking beads (Invitrogen, Eugene, USA). Only the blue laser was used for excitation of the stains. A 530/30 nm band-pass filter was used for the detection of green fluorescence (BL1) and a 695/40 nm band-pass filter was used for the detection of red fluorescence (BL3). A 488/10 band-pass filter was used for the detection of forward scatter (FSC) and side scatter (SSC). The flow rate was set to 100 µL/min and stop conditions were set to 100 µL of sample analyzed. The threshold (set on green fluorescence, BL1) and PMT-voltages were determined by using control samples: an untreated sample, a heat-killed sample (microwaved at 700 W for 1 min), a filter-sterilized sample, sterile unbuffered minimal medium and sterile PBS.

#### FACS

After staining, the intact cell population was sorted using a FACS Melody (BD, Erembodegem, Belgium) cell sorter equipped with a blue (488 nm) and yellow-green (561 nm) laser and a 100 µm sort nozzle. The performance of the instrument was checked using CS&T RUO Beads (BD, Erembodegem, Belgium) and drop delay was checked using FACS Accudrop RUO Beads (BD, Erembodegem, Belgium). Only the blue laser was used for excitation of the stains. A 527/32 nm band-pass filter was used for the detection of green fluorescence (PMT1) and a 700/54 nm band-pass filter was used for the detection of red fluorescence (PMT2). A 488/15 band-pass filter was used for the detection of side scatter (SSC). The threshold (set on green fluorescence, PMT1) and PMT-voltages were determined using control samples: an untreated sample, a heat-killed sample (microwaved at 700 W for 1 min), a filter-sterilized sample, sterile unbuffered minimal medium and sterile PBS. The intact cell population was gated based on the primary fluorescent channels (PMT1 and PMT2) (Supplementary 4; Figure S1). The number of events to sort was set to 250 000 (Supplementary 4; Table S4). After sorting, samples were stored at -20°C up until DNA extraction.

### Ion chromatography for determination of organic acids

Ion chromatography was used for the determination of organic acids (C1-C5). Samples were filtered (0.2 µm) and diluted 10-fold in dH_2_O and stored at -20°C up until actual chromatographic analysis. After thawing at room temperature until no more ice was visible, samples were analyzed according to Roume et al. using a 930 Compact IC Flex (Metrohm AG, Herisau, Switzerland) system with in-line bicarbonate removal (MCS) [53].

### DNA extraction

#### Extraction of sorted samples

First, samples were thawed at room temperature until no more ice was visible. DNA extractions were performed using the DNAeasy PowerSoil Pro kit (Qiagen, Venlo, Netherlands). 500 µL of sample was transferred to PowerBead Pro Tubes containing 250 µL of Solution CD1. Samples were disrupted in a PowerLyzer (Qiagen, Venlo, Netherlands) for 5 consecutive times at 4000 rpm for 15 seconds with 45-second cooldown intervals. All subsequent steps were performed following the standard manufacturer’s instructions. Samples were then stored at -20 °C up until further analysis.

#### Extraction of raw saliva and post-incubation samples

DNA extraction was performed using the DNAeasy PowerSoil Pro kit (Qiagen, Venlo, Netherlands). The cell pellet was mixed with 800 µL of Solution CD1 and transferred to a PowerBead Pro Tube with beads (provided with the kit). Next, samples were disrupted in a PowerLyzer (Qiagen, Venlo, Netherlands) for 5 consecutive times at 4000 rpm for 15 seconds with 45-second cooldown intervals. All subsequent steps were performed following the standard manufacturer’s instructions. After extraction, samples were stored at -20°C up until further analysis.

### Amplicon sequencing

For each sample, 15 µL genomic DNA extract (Supplementary 5) was sent to LGC genomics GmbH (Berlin, Germany) for library preparation and sequencing on an Illumina MiSeq platform with v3 chemistry with the primers 341F (5’ – CCT ACG GGN GGC WGC AG – 3’) and 785Rmod (5’ – GAC TAC HVG GGT ATC TAA KCC – 3’) [56].

### Data analysis

#### Analysis of flow cytometry data

Flow cytometry data were imported in R (version 4.2.0) [59] using the flowCore package (version 2.8.0) [60]. Data were transformed using the arcsine hyperbolic function [61], and gated manually on the primary fluorescent channels (BL1 and BL3) to remove background and to distinguish between intact and damaged cells (Supplementary 6). Total, intact and damaged cell concentrations were calculated based on the events in the cell gates and the volumetric measurements of the instrument.

Flow cytometric fingerprints (i.e. phenotypic fingerprints) were constructed by application of a Gaussian mixture mask to identify clusters within the flow cytometry data and were performed using the ‘*PhenoGMM*’ function of the Phenoflow package (version 1.1.2) [62]. To be able to generate the mask, for each timepoint (before incubation and after incubation), samples were subsampled to an equal number of cells so that every mouthwash treatment and every donor would be represented equally. This was done to avoid biased model training towards a specific donor or mouthwash treatment. The Gaussian mixture model (GMM) was optimized using the Bayesian information criterion (BIC) [63,64]. The GMM results in a one-dimensional (1D) vector for each sample that represents the number of cells allocated to each cluster in the model and can be regarded as a flow cytometric fingerprint (Supplementary 7). The parameters upon which the model was build were FSC-H, FSC-A, SSC-H, SSC-A, BL1-H, BL1-A, BL3-H, and BL3-A because these are expected to contain the most information [65]. Model output was first converted to relative abundances before further analysis. Converted model output was then used to perform Principal Coordinates Analysis (PCoA) based on Bray-Curtis dissimilarity (vegan package version 2.6-2) [66]. Statistical differences between mouthwash treatments were determined through distribution-independent analysis of similarities (ANOSIM) (vegan package version 2.6-2) [66]. P-values for pairwise comparisons were adjusted using the Benjamini and Hochberg method (stats package version 4.5.0) [59].

A random forest classifier to predict the most effective mouthwash (i.e. recommended mouthwash) for the saliva donor was trained on the *PhemoGMM* output of the intact microbial population of the control (PBS) before incubation using the caret package (version 6.0-92) [67]. The Matthews correlation coefficient (MCC) was used as a performance metric to optimize the classifier and was implemented using the mltools package (version 0.3.5) [68]. The MCC was used to account for class imbalance. The number of trees in the model was set to 500 and the model was trained using a three times repeated fivefold cross-validation scheme.

#### Analysis of organic acids

Organic acids were analyzed on the level of the molar concentration, as well as on the level of the cell concentration-normalized molar concentration. The normalized concentrations were calculated by dividing the molar concentration by the total cell concentration obtained through flow cytometry. Statistical differences between treatment groups were assessed using repeated measures analysis of variance (ANOVA) with Benjamini and Hochberg correction for multiple testing in the case of the molar concentration and using mixed-effects models for the normalized concentrations. The mixed-effects models were implemented using the lmerTest package (version 3.1-3) and pairwise comparisons with Benjamini and Hochberg correction were made using the emmeans package (version 1.9.0) [57,58]. Response to treatment was defined by assessing the relative reduction of organic acid concentration compared to the control (PBS). Strong responders were defined by a reduction larger than 60%, intermediate responders by a reduction between 20-60% and weak responders by a reduction lower than 20%. Additionally, the mouthwash that resulted in the highest reduction of organic acids was defined as the recommended mouthwash for the donor.

#### Processing of amplicon reads

Amplicon sequences were analyzed using the DADA2 package (version 1.30.0) in R as described by Callahan et al. [69]. After filtering, the average number of reads per sample was 30144 (Supplementary 5). ASVs were classified using the SILVA v138 training set [70]. The resulting read count table with taxonomic annotation was further processed in R (version 4.2.0). Rarefaction curves of all samples were used to assess if sufficient sequencing depth was obtained (vegan package version 2.6-2) [66] (Supplementary 8). Samples with a read count lower than 1300 were removed from further analysis. Quantitative microbial community composition was determined by multiplying the proportional read count data with microbial cell concentrations (flowCore package version 2.8.0) [60,71].

Alpha diversity was calculated using the iNEXT package (version 3.0.0) [72]. PCoA based on Bray-Curtis dissimilarity was performed on both the proportional and quantitative community composition using the vegan package (version 2.6-2) [66] and statistical differences between groups were determined through permutational analysis of variance (PERMANOVA) with Benjamini-Hochberg correction for multiple testing or through distribution-independent analysis of similarities (ANOSIM) with Benjamini-Hochberg correction for multiple testing (vegan package version 2.6-2) [66]. Distance-based Redundancy Analysis (db-RDA) based on Bray-Curtis dissimilarity was achieved using the vegan package (version 2.6-2) [66]. Results were visualized in a type 2 scaling correlation triplot and statistical significance of treatment, donor and timepoint (i.e. before and after incubation) were tested using permutation-based ANOVA for each term in the model. Due to multicollinearity between treatment and timepoint, a new model was built combining the two factors as one (further referred to as ‘Treatment’).

Differential abundance analysis on proportional data was performed using DESeq2 (version 1.42.0) [73]. For the quantitative community data mixed-effects models were used to account for repeated-measures for some donors. The Benjamini-Hochberg method was used for correcting for multiple testing. The mixed-effects models were implemented using the lmerTest package (version 3.1-3) [57].

Integrative analysis of the organic acid concentrations and the quantitative community composition was achieved by sparse Partial Least Squares (sPLS) analysis using the mixOmics package (6.26.0) [74]. First, a basic model was created in regression mode where the organic acid concentration was attempted to be explained by the quantitative community structure on the genus level. This was followed by model tuning, where the optimal number of variables and components were extracted in regression mode with correlation evaluation metrics (comp1 = 5, comp2 = 40 and comp3 = 25 for the quantitative community structure; comp1 = 1, comp2 =1 and comp3 = 4 for the organic acid concentration; n = 3 for the number of components). These parameters were used to construct a final sPLS model, which was in turn used to construct relevance network graphs to visualize structure associations.

## Results

### Changes in salivary microbiome composition after mouthwash treatment

To assess the impact of mouthwashes containing different fluoride formulations on the salivary microbiome composition, saliva samples of 12 different donors were treated with three distinct mouthwashes, namely Listerine Anti Caries, Elmex Anti Caries and Meridol. Analysis of cell concentrations revealed that Elmex and Meridol led to a higher reduction in intact cell concentration compared to Listerine and that the magnitude of the effect was donor dependent (Supplementary 9).

**Figure 2.**
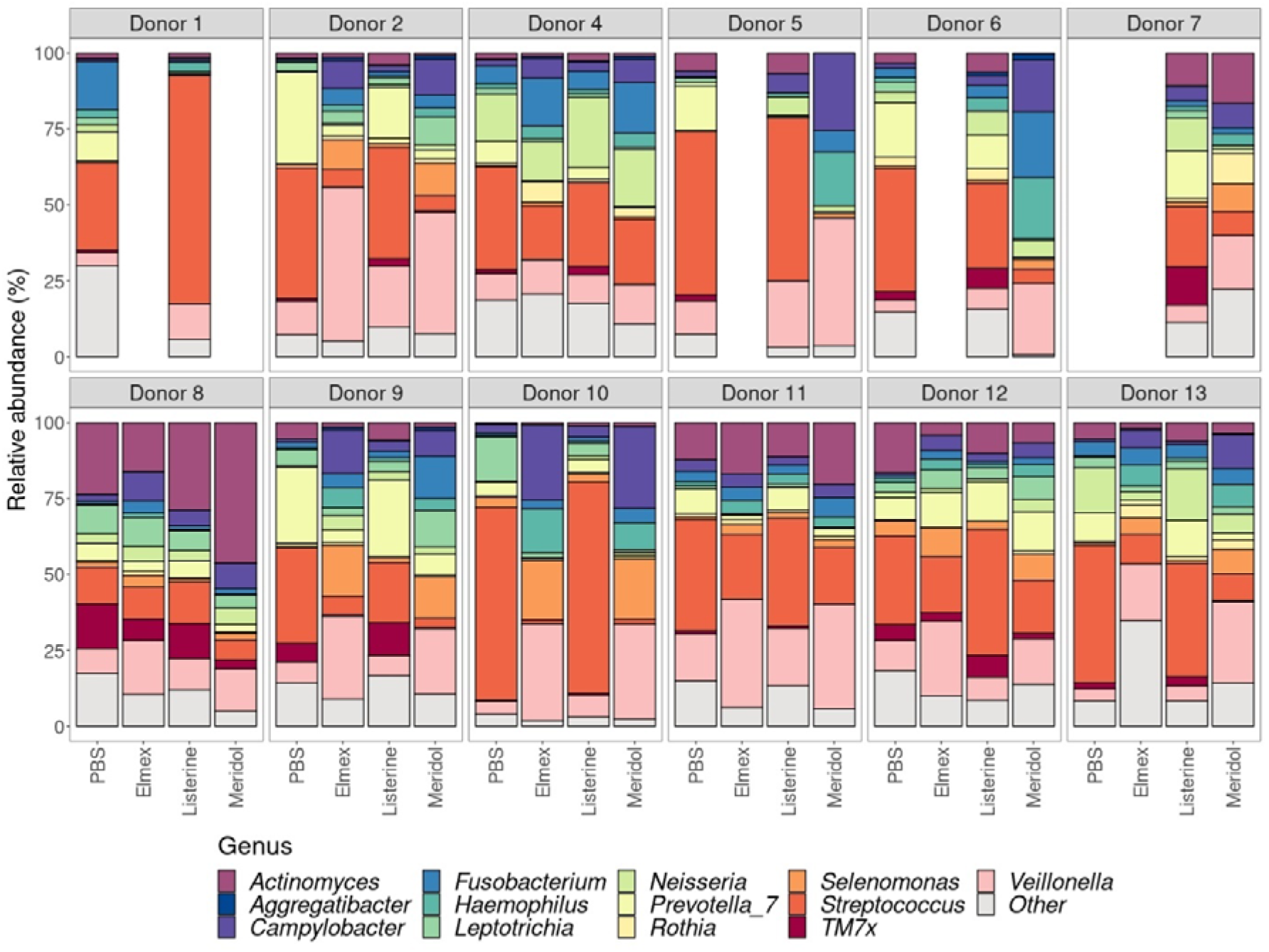
Relative abundances of the 13 most abundant bacterial genera in the intact cell population after mouthwash treatment. Lower abundant genera were grouped into ‘Other’. The missing samples are samples that were omitted from analysis due to a low number of read counts for these samples. Donor 3 is not included in the figure because DNA extraction failed for this donor.

The intact microbial populations after treatment was studied using 16S rRNA gene amplicon sequencing. Differences in community composition were observed between treatments on both the level of the proportional and quantitative community composition (Figure 2, Supplementary 10). No significant differences in alpha diversity could be observed between the treatments (Supplementary 11). There was a reduction in species richness (H0), Shannon diversity (H1) and inverse Simpson diversity (H2) for some donors (Donor 1, Donor 2, Donor 4, Donor 8, Donor 11 and Donor 12) after treatment with mouthwashes, however, this was not observed for all donors (Supplementary 11). PCoA based on Bray-Curtis dissimilarity showed grouping according to treatment for both the proportional and quantitative community data (Figure 3). Statistical analysis indicated significant differences in both proportional and quantitative community composition between PBS and Elmex (proportional (ANOSIM): *R* = 0.58, *P* < 0.001; quantitative (PERMANOVA): *F* = 4.95, *P* < 0.001), PBS and Meridol (proportional (ANOSIM): *R* = 0.50, *P* < 0.001; quantitative (PERMANOVA): *F* = 5.20, *P* < 0.001), Listerine and Elmex (proportional (ANOSIM): *R* = 0.51, *P* < 0.001; quantitative (PERMANOVA): *F* = 3.76, *P* < 0.001), and Listerine and Meridol (proportional (ANOSIM): *R* = 0.45, *P* < 0.001; quantitative (PERMANOVA): *F* = 3.95, *P* < 0.001). Similar patterns were observed in the flow cytometric fingerprint of the intact cell population after mouthwash treatment (Supplementary 12). Additionally, db-RDA revealed that all model factors (i.e. ‘Treatment’ and ‘Donor’) contributed significantly to the model for the proportional and quantitative community composition (Supplementary 13). Specifically, for the proportional microbiome composition, ‘Treatment’ contributed for 36.7% of the variation, while ‘Donor’ contributed for 31.9%. For the quantitative microbiome composition this was 37.2% for ‘Treatment’ and 23.4% for ‘Donor’.

**Figure 3.**
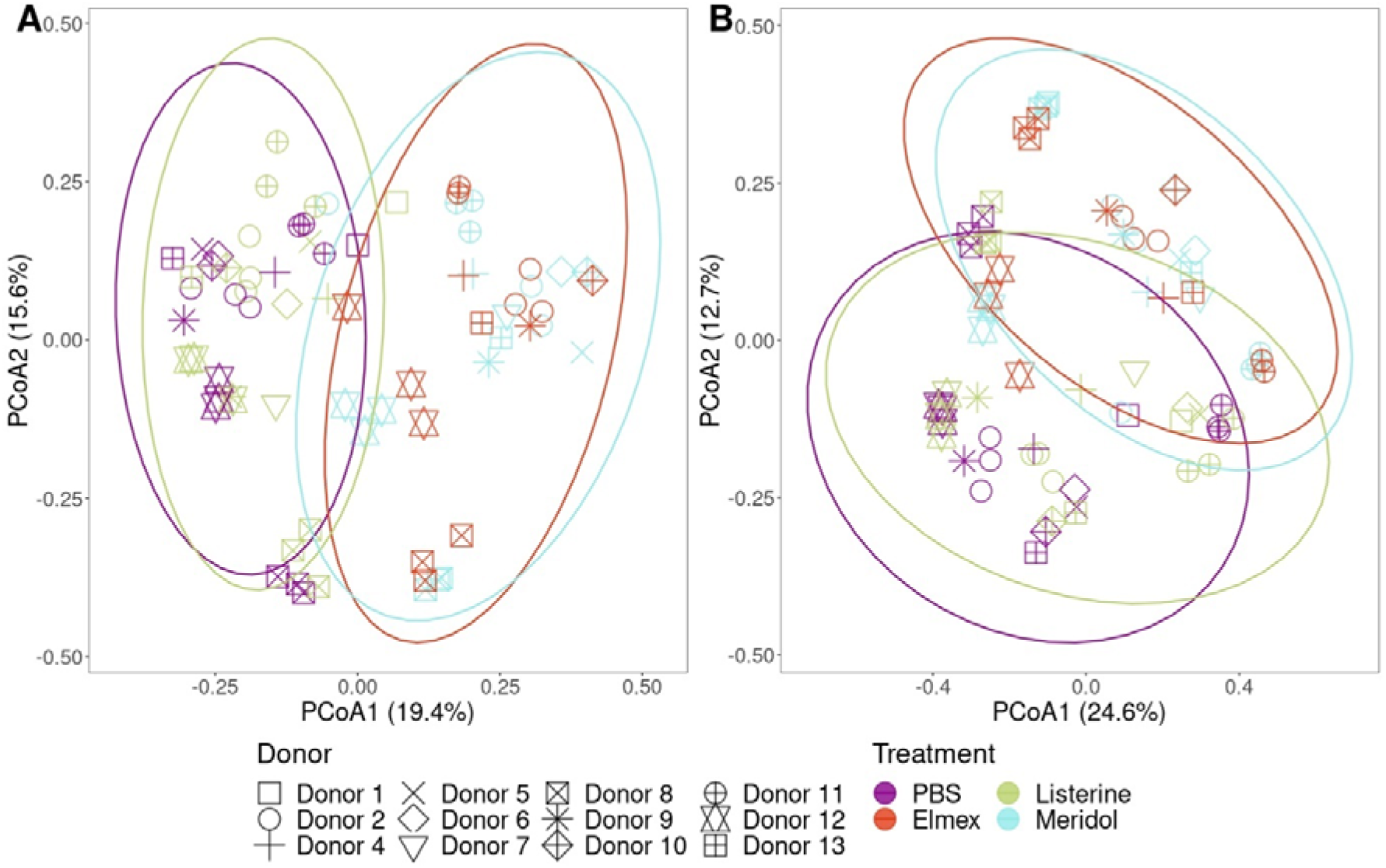
PCoA based on Bray-Curtis dissimilarity of the proportional (A) and quantitative (B) community composition of the intact cell population after mouthwash treatment (ASV level). Ellipses were drawn at the 95% confidence level. Donor 3 is not included in the figure because DNA extraction failed for this donor.

To determine how bacterial taxa were affected by the different mouthwashes, differentially abundant genera were identified for the proportional (Figure 4) and quantitative (Supplementary 14) community composition. Overall, Elmex and Meridol significantly reduced the abundance of more different bacterial genera compared to Listerine. For Elmex, ten genera were significantly reduced in the proportional community data, with an additional seven genera reduced in the quantitative data, totaling 17 genera. For Meridol, 19 genera were significantly reduced in the proportional community composition, of which 17 were also significantly reduced in the quantitative community data. In contrast, Listerine reduced only five genera in the proportional data, accompanied by nine additional genera in the quantitative data. Notably, no genera were uniquely reduced by Listerine or Elmex across both datasets, while Meridol uniquely reduced eight genera (*Megasphaera*, *Actinomyces*, *Pseudomonas*, *Pseudopropionibacterium*, *Prevotella*, *Alcaligenes*, *Sphingomonas* and *Atopobium*) in the proportional data, though no unique reductions were observed in the quantitative data. Six genera (*Streptococcus*, *Prevotella 7*, *Porphyromonas*, *TM7x*, *Butyrivibrio* and *Oribacterium*) in the proportional data and three genera (*Megasphaera*, *Oribacterium* and *TM7x*) in the quantitative data were reduced by both Elmex and Meridol, but not by Listerine. Additionally, *Leptotrichia* was reduced in the proportional community composition by both Listerine and Meridol, but not by Elmex. In terms of significantly increased genera, there were six genera in the proportional community composition and three genera in the quantitative community composition that were increased for Elmex, and one genus in the quantitative community composition that was increased for Meridol. This last genus (*Campylobacter*) was shared with Elmex.

**Figure 4.**
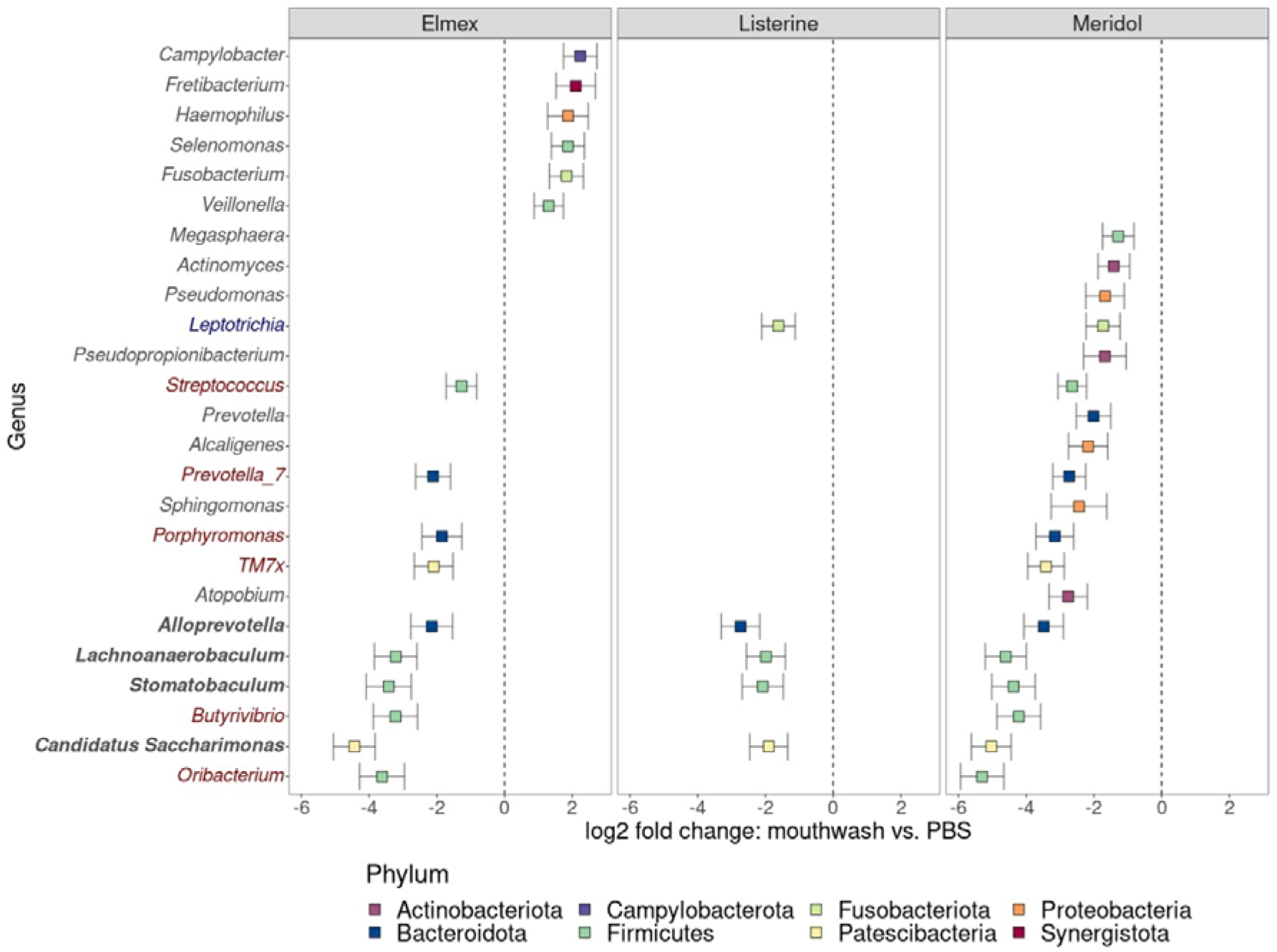
Differentially abundant genera for the different mouthwashes compared to the control (PBS) for the proportional community data. Only genera with a relative abundance higher than 1% in at least one of the samples are represented in the figure. Genus names in bold indicate that the genus was differentially abundant for all three mouthwashes, genus names in blue indicate genera that were differentially abundant for both Listerine and Meridol, and genus names in red indicate differentially abundant genera for both Elmex and Meridol. Error bars show the standard error for each genus and mouthwash.

### Anticaries potential of the mouthwashes

To determine the anticaries potential of the mouthwashes, treated saliva samples were incubated in minimal medium with sucrose as the sole carbon source so that the production of organic acids by the remaining microbial population could be assessed. Furthermore, the microbial community composition was studied to be able to link specific bacterial genera to the production of organic acids.

**Figure 5.**
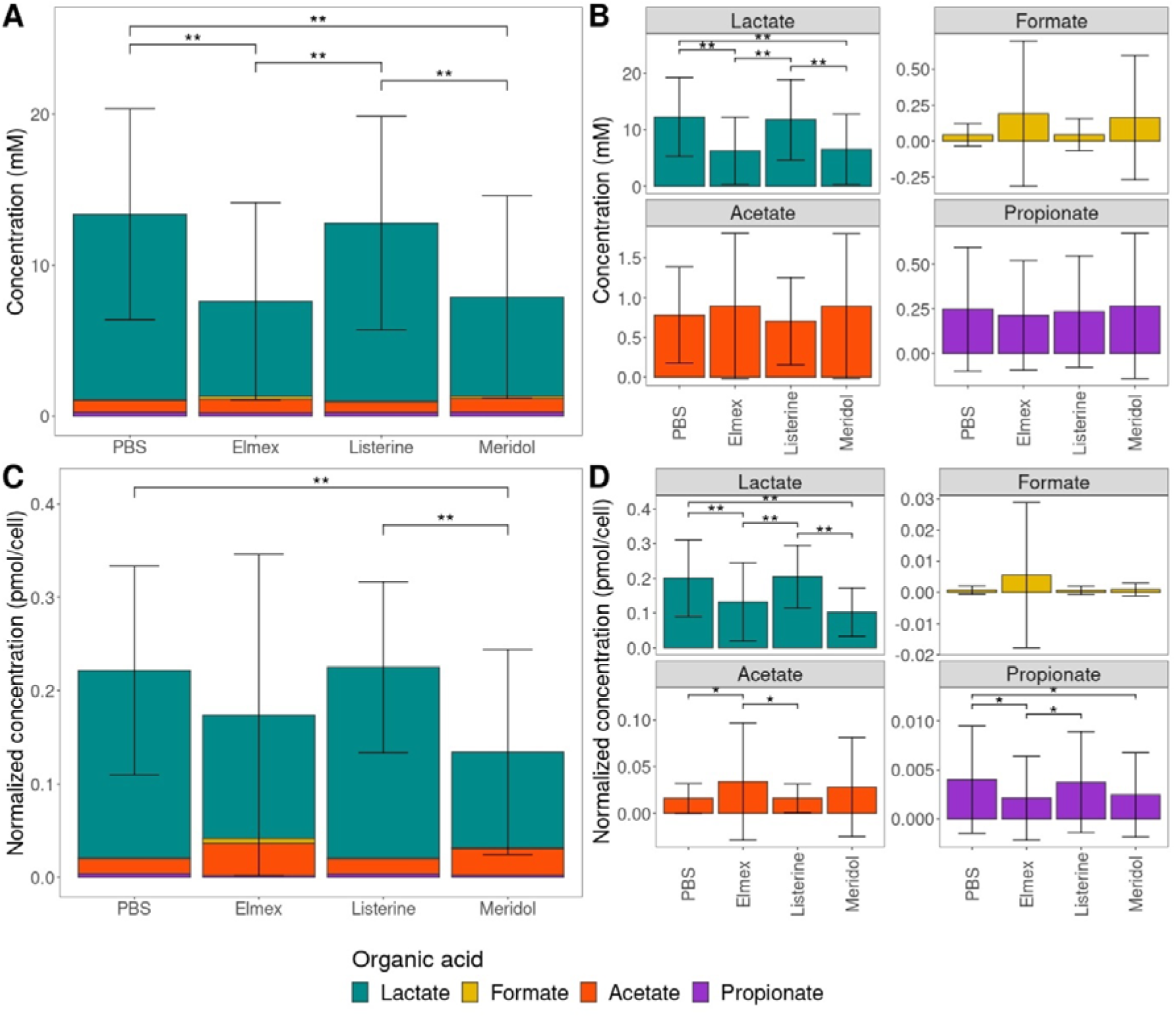
Comparisons of organic acids between treatments after incubation in sucrose-rich medium. The molar cell concentrations are shown in (A) and (B), and the total cell concentration-normalized acid concentrations are depicted in (C) and (D). Statistical differences were assessed using repeated measures ANOVA with Benjamini-Hochberg correction for multiple testing for the molar concentrations and mixed-effect models with Benjamini-Hochberg correction for multiple testing for the normalized acid concentrations. Statistical differences are indicated with asterisks (*P* < 0.05: *; *P* < 0.01: **). Error bars depict the standard deviation of the total acid concentration in (A) and (C) and for each acid in (B) and (D).

We found that Elmex and Meridol led to a significant decrease in total organic acid concentration (Figure 5A) and pH (Figure 6) compared to the control (PBS). This was not observed for Listerine.

Lactate was found to be the dominant organic acid for all treatments. When the organic acid concentration normalized on the total cell concentration was considered, only a significant decrease for Meridol was found compared to the control (Figure 5C). For the individual acids, Elmex and Meridol led to a reduction of acetate for both the molar concentration (Figure 5B) and the normalized concentration compared to the control (Figure 5D). In the normalized concentrations, there was also a reduction of propionate for Elmex and Meridol and an increase in acetate for Elmex compared to the control (Figure 5D). Moreover, large differences were observed between donors in both the molar acid concentration and normalized acid concentration (Supplementary 15).

**Figure 6.**
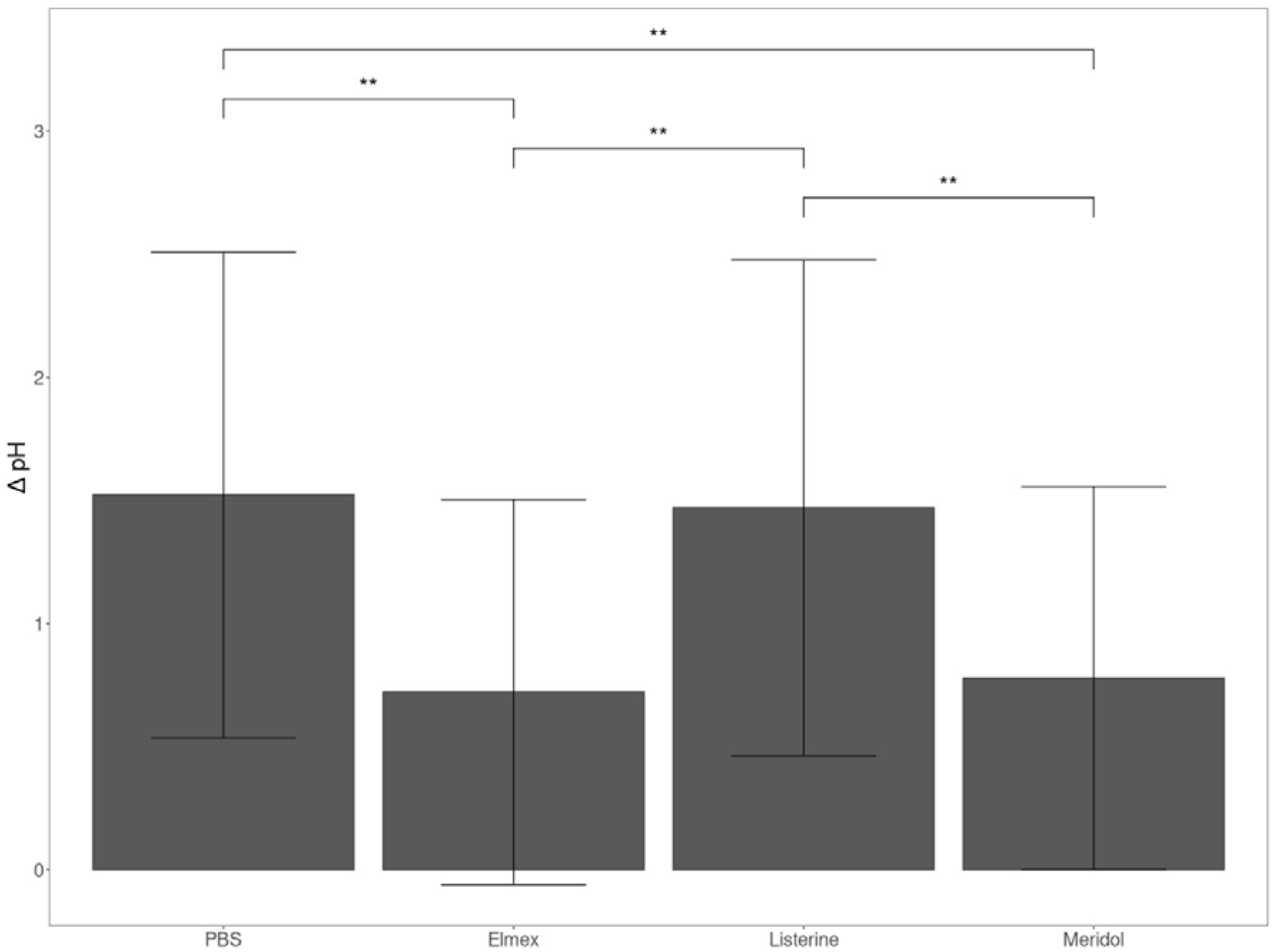
Drop in pH over the microaerophilic incubation of mouthwash-treated saliva samples in minimal medium with sucrose as carbon source for each mouthwash. Error bars show the standard deviation. Statistical differences were assessed using mixed-effect models with Benjamini-Hochberg correction for multiple testing and are indicated with asterisks (*P* < 0.05: *; *P* < 0.01: **).

The community composition after incubation showed that the communities were dominated by *Streptococcus* and that this dominance was more outspoken in the control (PBS) and Listerine samples (Supplementary 16). Differential abundance analysis (Supplementary 17) revealed that Elmex had 3 reduced genera (*Streptococcus*, *Prevotella 7* and *Alloprevotella*), Listerine had 2 reduced genera (*Leptotrichia* and *Alloprevotella*), and Meridol had 5 reduced genera (*Neisseria*, *Streptococcus*, *Prevotella7*, *Leptotrichia* and *Alloprevotella*) compared to the control for the proportional community composition. For the quantitative community composition only 2 genera (*Alloprevotella* and *Leptotrichia*) were reduced compared to the control and shared between all mouthwashes. All genera that were found to be differentially abundant in the quantitative community data, were also found to be different in the proportional community data, except for *Leptotrichia* with Elmex, which was only found to be differentially abundant in the quantitative community data.

To be able to link the organic acids to specific genera, a relevance network was constructed based on sparse Partial Least Squares (sPLS) analysis using the molar acid concentrations and quantitative community structure (Figure 7). We found intermediate positive correlation (0.61) between lactate and *Streptococcus*. Additionally, we found an intermediate to high positive correlation between acetate and *Prevotella 7* (0.57), *Veillonella* and acetate (0.65), propionate and *Prevotella 7* (0.67), propionate and *Veillonella* (0.77), and propionate and *Prevotella* (0.51).

**Figure 7.**
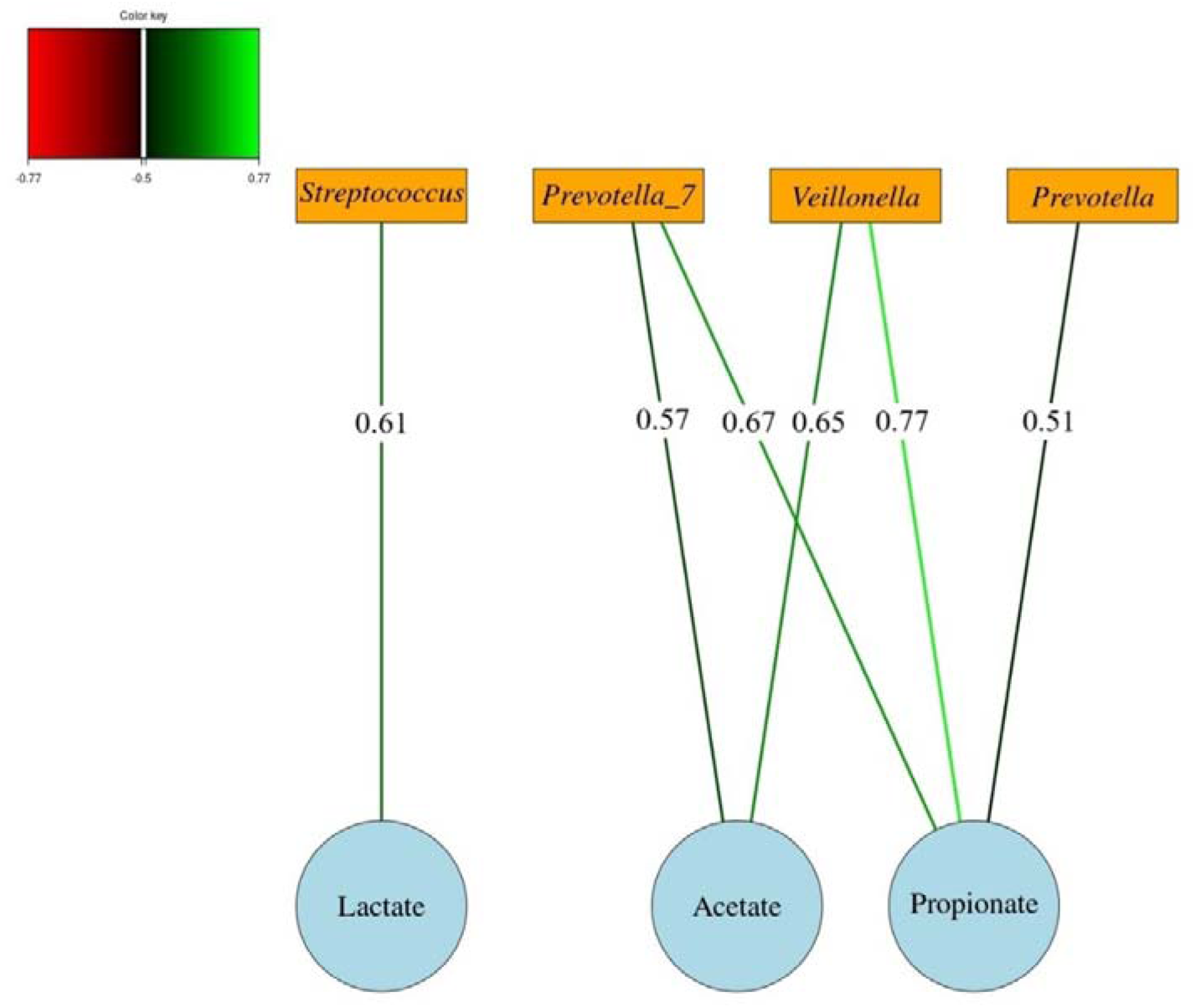
Relevance network for the sPLS regression of the molar organic acid concentration and absolute genus abundance of mouthwash-treated saliva after incubation in minimal medium with sucrose as carbon source. Only correlations between acids and genera with an absolute value greater than 0.50 are shown in the network. Positive correlations are shown by the green lines.

### Response to treatment by assessment of the genotypic and phenotypic fingerprint

Since high interindividual variability was observed in the anticaries potential of the different mouthwashes (Supplementary 15), we assessed if the response to treatment of a donor could be captured in the phenotypic (i.e. flow cytometric) and genotypic (i.e. proportional and quantitative community composition) fingerprint of the initial intact salivary microbiota. Therefore, the fingerprints of the different donors were labelled based on which mouthwash resulted in the highest reduction of molar concentration of organic acids (i.e. the recommended mouthwash). All donors were assigned to either Elmex or Meridol, as treatment with Listerine never resulted in the highest reduction in organic acids. PCoA based on Bray-Curtis dissimilarities revealed that only separate groups could be distinguished in the flow cytometric fingerprint and not in the genotypic fingerprints of the microbial community (Figure 8). This was confirmed by statistical analysis where no differences between recommended mouthwash groups were found for the proportional and quantitative community composition, while significant differences were found between recommended mouthwash groups in the flow cytometric fingerprints (ANOSIM: *R* = 0.20, *P* = 0.01).

**Figure 8.**
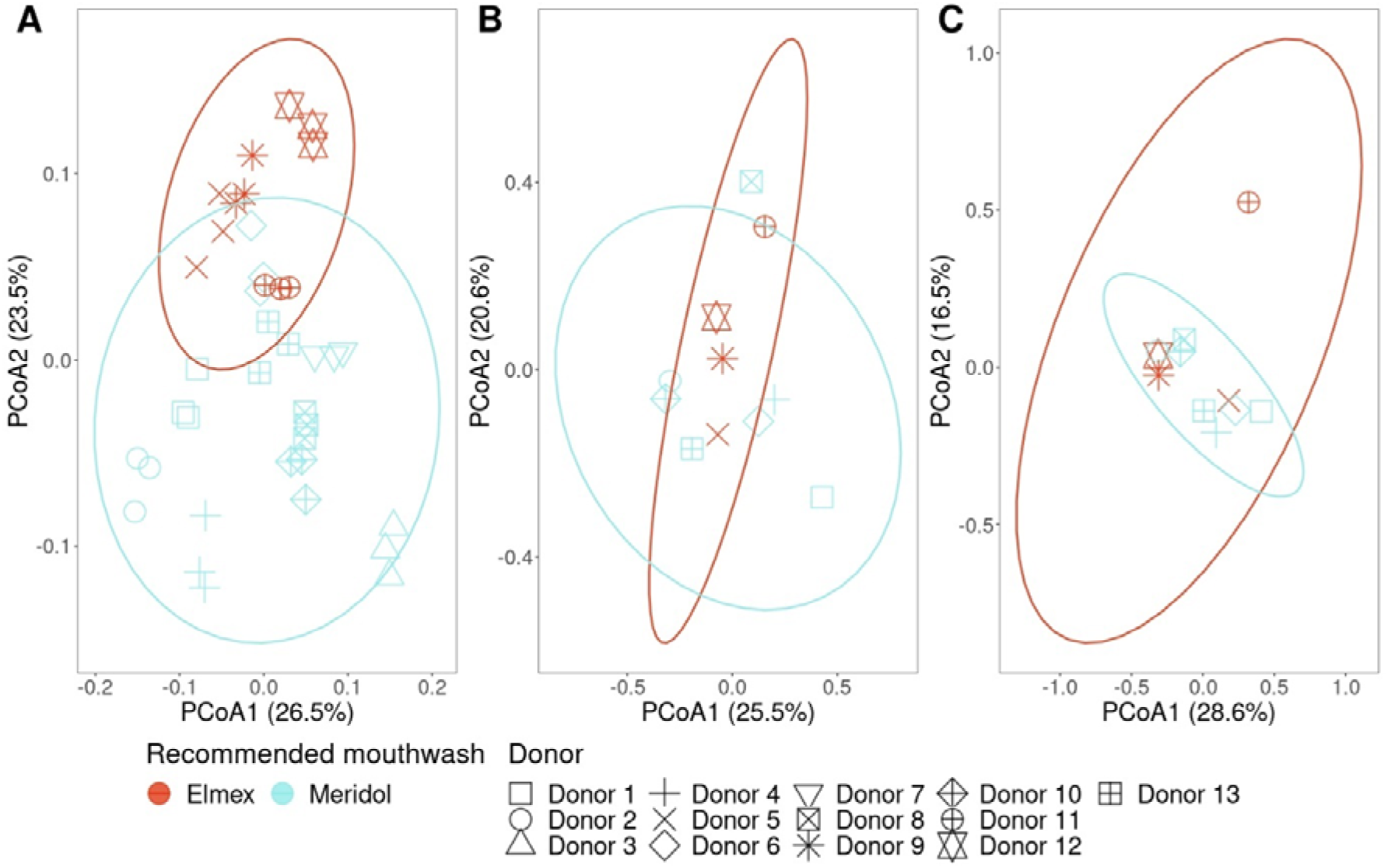
PCoA of the intact cell population of the control (PBS) before incubation based on Bray-Curtis dissimilarity for the flow cytometric fingerprint (A), proportional community composition (B), and quantitative community composition (C). Colors show the recommended mouthwash for each donor which was determined based on the mouthwash that resulted in the highest reduction in total molar organic acid concentration compared to the control. Ellipses were drawn at the 95% confidence level. Donor 3 is not included in (B) and (C) because DNA extraction failed for this donor.

Because statistical differences were found between recommended mouthwash groups in the flow cytometric fingerprints, these fingerprints were used to train a random forest classifier for the prediction of the recommended mouthwash of saliva donors (Figure 9). The classifier yielded an MCC of 0.95. The class accuracy for Elmex was 100% and 95.6% for Meridol.

**Figure 9.**
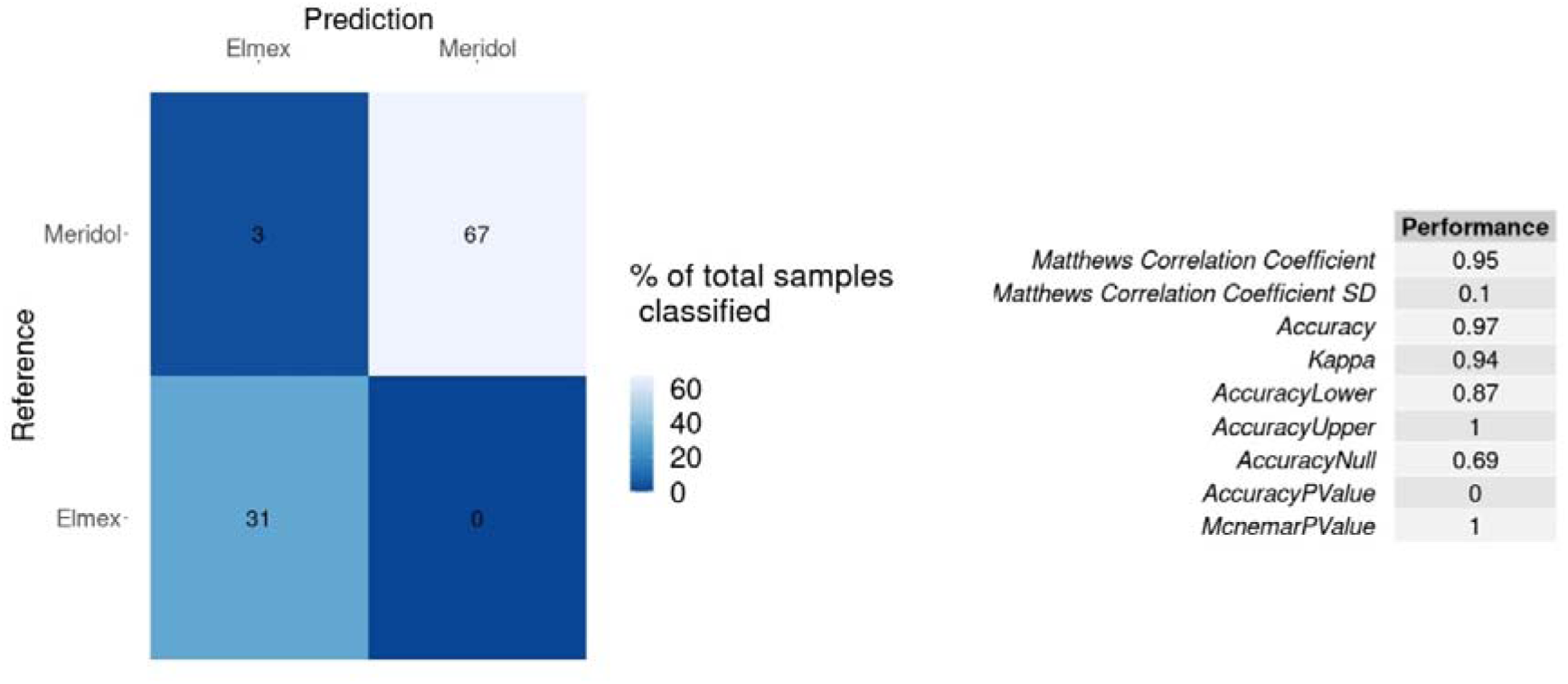
Confusion matrix and performance metrics for the random forest classifier trained to predict the recommended mouthwash for saliva donors. The model was trained using a three-times repeated fivefold cross-validation scheme.

## Discussion

Different formulations of fluoride are used to prevent dental caries and are reported to have differing efficacy against oral bacteria [15,18,31,34,37]. Additionally, large interindividual differences in oral microbiome composition exist and differing responses to treatment are to be expected [40–42]. Because there is no existing literature that compares changes in oral microbiome composition between the different formulations of fluoride, we studied changes in salivary microbiome composition after treatment with different fluoride-containing mouthwashes. Moreover, we investigated the anticaries potential of the mouthwashes by assessment of organic acids produced by the remaining microbial community after treatment and evaluated interindividual differences in response to treatment between saliva donors. Our results showed that a shift in salivary microbiome composition occurred and that this shift was dependent on the mouthwash used. Furthermore, we found that the anticaries potential was also dependent on the mouthwash and observed interindividual differences in anticaries potential between donors. Finally, we could capture the response to treatment in the flow cytometric fingerprint of the initial intact cell population of the donor and successfully train a random forest classifier predicting the most effective mouthwash.

### Change in salivary microbiome composition depends on the mouthwash

By assessing the salivary microbiome composition, we found that the different mouthwashes induced a taxonomic shift in the intact salivary microbiota (Figure 3) and found differences in differentially abundant genera for the different mouthwashes (Figure 4, Supplementary 14). The effect of Elmex and Meridol was comparable, however, treatment with Listerine resulted in a community that was more like the control (PBS). Meridol and Elmex effectively reduced *Oribacterium* and *TM7x* in both the proportional and quantitative community composition, *Megasphaera* in the quantitative community composition, and *Streptococcus*, *Prevotella 7*, *Porphyromonas* and *Butyrivibrio* in the proportional community composition, whereas Listerine did not. All these genera have been implicated in dental caries, except for *Porphyromonas* which plays an important role in periodontal disease [6,11,13,14,75–78]. Additionally, Elmex and Meridol damaged a larger proportion of the salivary microbiota (Supplementary 9). Thus, these mouthwashes are more effective in reducing oral bacteria, and by extension oral pathogens. This can be explained by the different (active) ingredients in the mouthwashes. Listerine only contains NaF as an active fluoride ingredient, while Elmex contains both NaF and AmF, and Meridol contains AmF and SnF_2_. Simultaneously, Listerine contains essential oils with demonstrated antimicrobial properties and other ingredients with antimicrobial effects (e.g. sodium benzoate) are present in the different mouthwashes as well [79,80]. Previous research showed that NaF was less effective in reducing pathogenic oral bacteria compared to AmF and SnF_2_, illustrating the importance of the antimicrobial properties of the fluoride counter ions, rather than the antimicrobial effect of fluoride [31,34,37]. Furthermore, the antimicrobial effect of fluoride is expected to be limited, as the final concentration in our experiment is low and the importance of the antimicrobial properties of fluoride in caries prevention is questioned [23,39]. In our data, the added effect of SnF_2_ in the mouthwash seemed to have a limited effect when AmF was already present. Only in the proportional microbiome composition more taxa for Meridol than for Elmex were found to be reduced compared to the control (Figure 4). In addition, the community composition between Elmex and Meridol was similar (Figure 3). Literature reports that the antimicrobial effects of SnF_2_ are comparable with AmF [38]. Simultaneously, Sn^2+^ has been suggested to exhibit selective binding towards specific pathogenic species [30]. Since the total concentration of fluorides in the two mouthwashes (Elmex and Meridol) was the same and because AmF was shared between the two mouthwashes, combined with the presence of other antimicrobial ingredients in the different mouthwashes, we argue that the overall antimicrobial effect of the mouthwash is the most important driver for the change in community structure. Thus, also the other (active) ingredients in the mouthwashes should be considered. Comparing pure NaF, AmF and SnF_2_ to each other may shed more light on the specificity of the fluoride compounds towards certain bacterial strains.

Remarkably, there were also genera that were increased after mouthwash treatment. Since the mouthwashes contained antimicrobial compounds and no incubation was performed before analysis, these increased genera should be considered biased. There were more increased genera in the proportional community data, which illustrates the possible bias that can be introduced when resorting to differential abundance analysis methods for estimating differentially abundant taxa based on proportional data [81]. All increased genera in both the proportional and quantitative microbiome composition are all normal constituents of the oral microbiota and therefore should not be seen as being increased due to contamination. However, bias introduced in the methodology could be the cause of this phenomenon. For example, it could be that staining efficiencies of these specific taxa were different under the different chemical conditions induced by the different mouthwashes when using FACS for retaining the intact cell population for further analysis [82].

### Anticaries potential is dependent on the mouthwash

Through assessment of organic acids after incubation in minimal medium with sucrose, we found that the anticaries potential of Elmex and Meridol was greater than that of Listerine. When lower amounts of acids were produced, the anticaries potential was deemed greater. More specifically, Elmex and Meridol led to a reduction of total molar organic acid concentration compared to the control, while Listerine did not. For Meridol, a significant reduction was only found when the organic acid concentration was normalized on the total cell concentration in the sample. Although not significant, a reduction was also observed for Elmex in the normalized total acid concentration. The better anticaries potential of Elmex and Meridol was supported by the observed pH-drop over the incubation, where Elmex and Meridol had a significantly smaller pH-drop compared to both the control and Listerine. We argue that the differences between the mouthwashes are the result of a different microbial community composition at the start of the incubation between the mouthwashes. We showed that the mouthwashes exerted a different effect on the intact microbiome composition, thus different species were present for regrowth after mouthwash treatment. Given Meridol was the only mouthwash that resulted in a significant reduction in the total normalized acid concentration, it led to the most favorable microbiological shift for treatment of dental caries on the level of the entire donor population. Moreover, the molar concentration and normalized concentration of lactate were significantly reduced for both Elmex and Meridol, but not for Listerine. Since lactate is the most abundant organic acid produced in our experiment, we argue that the difference in total acid concentration is mostly driven by the difference in concentration of lactate. Network analysis revealed a positive correlation (0.61) between *Streptococcus* and lactate (Figure 7). This finding is consistent with literature, where lactate has been reported to be the predominant end metabolite of oral Streptococci at low pH under anaerobic conditions [83,84]. Differential abundance analysis after incubation showed that *Streptococcus* was reduced for Elmex and Meridol in the proportional community composition and not for Listerine, compared to the control (Supplementary 17). Thus, the reduction in lactate for Elmex and Meridol is likely the result of the reduction in *Streptococcus*. Propionate was also found to be reduced for Elmex and Meridol compared to the control in the normalized acid concentrations (Figure 5). The relevance network showed that propionate was positively correlated with *Prevotella 7* (0.67), *Veillonella* (0.77) and *Prevotella* (0.51) (Figure 7). Indeed, members of *Prevotella* and *Veillonella* are known to produce propionate [85–87]. Differential abundance analysis after incubation showed that *Prevotella 7* was reduced for Elmex and Meridol compared to the control, although this was not the case for *Veillonella* and *Prevotella* (Supplementary 17). However, the relative abundance of *Prevotella 7* in the community was greater than that of *Prevotella* and *Veillonella* (Supplementary 16). Thus, the reduction in propionate may be driven by the reduction in *Prevotella 7* for Elmex and Meridol. Last, we found that acetate was increased for Elmex compared to the control in the normalized acid concentrations and that acetate was positively correlated with *Prevotella 7* (0.57) and *Veillonella* (0.65) (Figure 5; Figure 7). Again, members of these genera are known to produce acetate [85,86]. Nevertheless, these genera were not found to be significantly more prevalent for Elmex compared to the control, and *Prevotella 7* was even found to be reduced in the proportional community data (Supplementary 17). It is important to note that metabolic interactions in the oral cavity are common and that organic acids can be used as carbon sources by different oral bacteria [88]. Thus, the differential abundance of single species may not be enough to fully explain differences in specific organic acids.

### The response to treatment can be captured in the flow cytometric fingerprint

Based on the high interindividual differences between donors regarding response to treatment, a recommended mouthwash for each saliva donor was defined by choosing the mouthwash that led to the highest reduction in molar organic acid concentration compared to the control. This resulted in either Elmex or Meridol as the recommended mouthwash for our study cohort. We chose the molar organic acid concentration for our recommendation because it is more relevant than the normalized organic acid concentration in a clinical context. It is the absolute concentration of acids that drives tooth demineralization [6,89]. We were able to distinguish groups between donors depending on the recommended mouthwash (i.e. response to treatment) in the flow cytometric fingerprint but not in the genotypic fingerprint of the initial intact microbial community (Figure 8). Hence, the phenotypic (i.e. flow cytometric) fingerprint provided information on the response to treatment that could not be captured in the genotypic fingerprint. Previous research has shown that the phenotype of genetically identical microbial populations can play an important role in response to environmental changes [90,91]. Therefore, we argue that the phenotype of the salivary microbiome also influences the response to mouthwash treatment. Additionally, as the effects of Elmex and Meridol were similar regarding shifts in microbial community composition for our data, it may be that higher taxonomic resolution would lead to better clustering of the recommended mouthwash groups in the genotypic fingerprints.

We successfully trained a random forest classifier predicting the recommended mouthwash for each donor based on the flow cytometric fingerprint (Figure 9). This result shows the potential of using flow cytometry for predictive modelling for tailoring treatments to patients. To increase accuracy in determining the most suitable treatment for patients, a combination of an *in vitro* assay, a genotypic fingerprint and a flow cytometric fingerprint could be considered as well. It should be noted that a limitation of our study is that we determined the anticaries potential of the mouthwashes by an *in vitro* incubation. To confirm our findings, *in vivo* tests should be performed as well. Furthermore, the number of participants in this study is relatively small. If the workflow is to be applied in clinical practice, a more extensive study population must be considered.

## Conclusion

To conclude, we found that a shift in the salivary microbiome composition occurred after treatment with fluoride-containing mouthwashes and that this shift was dependent on the formulation of the mouthwash. The shift was least outspoken for Listerine (NaF), while the more prominent effects of Elmex (AmF and NaF) and Meridol (AmF and SnF_2_) were similar. Meridol seemed to affect more bacterial taxa and had the best anticaries potential on the level of the study population. On the level of the individual saliva donor, either Meridol or Elmex showed the best anticaries potential. Last, we were able to capture the response to treatment in the flow cytometric fingerprint of the initial intact salivary microbiome and could successfully train a classifier predicting the most suitable mouthwash for the saliva donor using this fingerprint.

## Code and data availability

Raw flow cytometry data and metadata are available at FlowRepository under ID FR-FCM-Z874. Primer clipped sequence reads are available at NCBI as Sequence Read Archive (SRA) under BioProject ID PRJNA1155198 (https://www.ncbi.nlm.nih.gov/sra/PRJNA1155198). The data analysis scripts are available at https://github.com/famermans/MouthwashCaries_SalivaryMicrobome.

## Acknowledgements

The authors would like to thank Tim Lacoere for providing the figure of the experimental workflow, and Yorick Minnebo, Marc De Doncker and Valérie Mattelin for critically reading the manuscript. This work was supported by Research Foundation – Flanders (FWO, Belgium) (grant number: G0B2719N).

## Author contributions

FM, NB and WT conceived and designed the study. FM and BP performed the experimental work and analyzed the data. MS performed the clinical evaluations of the donors. FM wrote the manuscript. FM created the figures. WT and NB supervised the findings of this work. All authors reviewed and approved the manuscript.

