## Supplementary information for "Interindividual differences in caries potential of the salivary microbiome after mouthwash exposure"

**Supplementary 1. Characteristics of saliva donors**

Table S1. Information on saliva donors. Plaque level, average probing pocket depth (PPD), clinical attachment loss (CAL), and bleeding on probing (BOP) were assessed by a dentist. When BOP was ≥ 10 %, donors were considered suffering from gingivitis.

| **Donor** | **Sex** | **Age** | **Ethnicity** | **Number of natural teeth** | **Plaque level (%)** | **Average PPD** | **CAL** | **BOP (%)** | **Gingivitis** |
| --- | --- | --- | --- | --- | --- | --- | --- | --- | --- |
| Donor 1 | Female | 29 | Caucasian | >= 20 | 78.61 | 2.47 | 3.00 | 23.12 | Yes |
| Donor 2 | Female | 26 | Caucasian | >= 20 | 64.74 | 2.23 | 3.30 | 8.09 | No |
| Donor 3 | Female | 33 | Caucasian | >= 20 | 60.12 | 2.20 | 2.63 | 8.33 | No |
| Donor 4 | Female | 56 | Caucasian | 10-19 | 86.88 | 2.56 | 4.33 | 10.00 | Yes |
| Donor 5 | Male | 30 | Caucasian | >= 20 | 86.90 | 2.44 | 4.33 | 2.38 | No |
| Donor 6 | Female | 33 | Caucasian | >= 20 | 74.40 | 2.45 | 4.25 | 17.26 | Yes |
| Donor 7 | Female | 29 | Caucasian | >= 20 | 15.2 | 2.32 | 4.25 | 12.28 | Yes |
| Donor 8 | Female | 35 | Brancos | >= 20 | 76.79 | 2.26 | 3.47 | 5.36 | No |
| Donor 9 | Male | 25 | Caucasian | >= 20 | 85.12 | 2.43 | 3.22 | 10.71 | Yes |
| Donor 10 | Female | 22 | Caucasian | >= 20 | 45.83 | 1.96 | 2.50 | 2.98 | No |
| Donor 11 | Male | 29 | Caucasian | >= 20 | 53.57 | 2.44 | 3.31 | 3.57 | No |
| Donor 12 | Female | 25 | Caucasian | >= 20 | 49.40 | 2.24 | 2.00 | 4.76 | No |
| Donor 13 | Male | 32 | Caucasian | >= 20 | 97.02 | 2.57 | 3.55 | 24.40 | Yes |

**Supplementary 2. Ingredient list of mouthwashes**

Table S2. Ingredient list of the different mouthwashes. Checkmarks indicate whether the ingredient is present in the mouthwash.

| **Ingredient** | **Listerine Anti Caries** | **Elmex Anti Caries** | **Meridol** |
| --- | --- | --- | --- |
| Aqua | ✔ | ✔ | ✔ |
| Aroma | ✔ | ✔ | ✔ |
| Benzoic acid | ✔ |  |  |
| Caffeine | ✔ |  |  |
| Camellia sinensis leaf extract | ✔ |  |  |
| CI 42051 |  |  | ✔ |
| CI 42053 | ✔ |  |  |
| CI 47005 | ✔ |  |  |
| Eucalyptol | ✔ |  |  |
| Glycerin |  | ✔ |  |
| Levulinic acid |  | ✔ |  |
| Menthol | ✔ |  |  |
| Methyl salicylate | ✔ |  |  |
| Olaflur |  | ✔ | ✔ |
| PEG-40 hydrogenated castoir oil |  | ✔ | ✔ |
| Poloxamer 407 | ✔ |  |  |
| Propylene glycol | ✔ | ✔ |  |
| PVP |  |  | ✔ |
| Saccharin |  | ✔ |  |
| Sodium anisate |  | ✔ |  |
| Sodium benzoate | ✔ | ✔ |  |
| Sodium fluoride | ✔ | ✔ |  |
| Sodium lauryl sulfate | ✔ |  |  |
| Sodium levulinate |  | ✔ |  |
| Sodium saccharin | ✔ |  | ✔ |
| Sorbitol | ✔ |  |  |
| Stannous fluoride |  |  | ✔ |
| Sucralose | ✔ |  |  |
| Thymol | ✔ |  |  |
| Xylitol |  |  | ✔ |

**Supplementary 3. Determination of pH**

Table S3. pH of the samples before and after incubation.

| **Donor** | **Treatment** | **Replicate** | **pH before incubation** | **pH after incubation** |
| --- | --- | --- | --- | --- |
| Donor 1 | PBS | 1 | 7.39 | 6.87 |
|  |  | 2 | 7.38 | 6.87 |
|  |  | 3 | 7.39 | 6.91 |
|  | Elmex | 1 | 7.38 | 6.99 |
|  |  | 2 | 7.38 | 6.99 |
|  |  | 3 | 7.39 | 6.99 |
|  | Listerine | 1 | 7.38 | 6.95 |
|  |  | 2 | 7.43 | 6.83 |
|  |  | 3 | 7.43 | 6.66 |
|  | Meridol | 1 | 7.42 | 6.99 |
|  |  | 2 | 7.43 | 7.04 |
|  |  | 3 | 7.39 | 7.04 |
| Donor 2 | PBS | 1 | 7.35 | 5.48 |
|  |  | 2 | 7.34 | 6.23 |
|  |  | 3 | 7.38 | 6.61 |
|  | Elmex | 1 | 7.38 | 6.61 |
|  |  | 2 | 7.48 | 7.04 |
|  |  | 3 | 7.48 | 6.91 |
|  | Listerine | 1 | 7.39 | 6.74 |
|  |  | 2 | 7.43 | 6.53 |
|  |  | 3 | 7.38 | 6.67 |
|  | Meridol | 1 | 7.43 | 6.85 |
|  |  | 2 | 7.38 | 6.83 |
|  |  | 3 | 7.38 | 6.74 |
| Donor 3 | PBS | 1 | 7.42 | 7.02 |
|  |  | 2 | 7.38 | 7.09 |
|  |  | 3 | 7.38 | 7.09 |
|  | Elmex | 1 | 7.38 | 7.15 |
|  |  | 2 | 7.33 | 7.09 |
|  |  | 3 | 7.38 | 7.14 |
|  | Listerine | 1 | 7.38 | 7.09 |
|  |  | 2 | 7.43 | 7.09 |
|  |  | 3 | 7.38 | 7.09 |
|  | Meridol | 1 | 7.38 | 7.09 |
|  |  | 2 | 7.43 | 7.14 |
|  |  | 3 | 7.38 | 7.19 |
| Donor 4 | PBS | 1 | 7.38 | 6.78 |
|  |  | 2 | 7.38 | 6.2 |
|  |  | 3 | 7.39 | 6.08 |
|  | Elmex | 1 | 7.38 | 6.75 |
|  |  | 2 | 7.38 | 6.79 |
|  |  | 3 | 7.38 | 6.75 |
|  | Listerine | 1 | 7.38 | 6.5 |
|  |  | 2 | 7.48 | 6.68 |
|  |  | 3 | 7.43 | 6.71 |
|  | Meridol | 1 | 7.44 | 6.87 |
|  |  | 2 | 7.43 | 6.86 |
|  |  | 3 | 7.38 | 6.85 |
| Donor 5 | PBS | 1 | 7.14 | 4.52 |
|  |  | 2 | 7.19 | 4.55 |
|  |  | 3 | 7.19 | 4.55 |
|  | Elmex | 1 | 7.19 | 7 |
|  |  | 2 | 7.14 | 6.92 |
|  |  | 3 | 7.19 | 6.75 |
|  | Listerine | 1 | 7.18 | 4.76 |
|  |  | 2 | 7.18 | 4.76 |
|  |  | 3 | 7.14 | 4.68 |
|  | Meridol | 1 | 7.14 | 6.95 |
|  |  | 2 | 7.19 | 5.88 |
|  |  | 3 | 7.14 | 6.5 |
| Donor 6 | PBS | 1 | 6.97 | 6.23 |
|  |  | 2 | 6.95 | 6.43 |
|  |  | 3 | 6.95 | 6.34 |
|  | Elmex | 1 | 6.99 | 6.69 |
|  |  | 2 | 6.99 | 6.69 |
|  |  | 3 | 6.95 | 6.6 |
|  | Listerine | 1 | 7.09 | 6.38 |
|  |  | 2 | 6.93 | 6.29 |
|  |  | 3 | 6.99 | 6.56 |
|  | Meridol | 1 | 6.99 | 6.56 |
|  |  | 2 | 6.99 | 6.73 |
|  |  | 3 | 6.99 | 6.64 |
| Donor 7 | PBS | 1 | 6.99 | 6.34 |
|  |  | 2 | 6.99 | 6.42 |
|  |  | 3 | 6.99 | 6.25 |
|  | Elmex | 1 | 6.99 | 6.65 |
|  |  | 2 | 6.98 | 6.72 |
|  |  | 3 | 7 | 6.64 |
|  | Listerine | 1 | 6.99 | 6.38 |
|  |  | 2 | 6.99 | 6.47 |
|  |  | 3 | 6.99 | 6.34 |
|  | Meridol | 1 | 7.04 | 6.72 |
|  |  | 2 | 6.99 | 6.62 |
|  |  | 3 | 6.99 | 6.64 |
| Donor 8 | PBS | 1 | 7.04 | 4.92 |
|  |  | 2 | 7.04 | 4.79 |
|  |  | 3 | 7.04 | 4.92 |
|  | Elmex | 1 | 7.04 | 5.17 |
|  |  | 2 | 7.03 | 5.21 |
|  |  | 3 | 7.04 | 5.22 |
|  | Listerine | 1 | 7.04 | 4.92 |
|  |  | 2 | 7.04 | 4.84 |
|  |  | 3 | 7.04 | 4.8 |
|  | Meridol | 1 | 7.04 | 5.1 |
|  |  | 2 | 6.99 | 5.32 |
|  |  | 3 | 7.04 | 5.14 |
| Donor 9 | PBS | 1 | 7.12 | 4.15 |
|  |  | 2 | 7.14 | 4.15 |
|  |  | 3 | 7.09 | 4.12 |
|  | Elmex | 1 | 6.92 | 5.1 |
|  |  | 2 | 7.06 | 5.1 |
|  |  | 3 | 7.11 | 6.36 |
|  | Listerine | 1 | 7.08 | 4.14 |
|  |  | 2 | 7.04 | 4.16 |
|  |  | 3 | 7.05 | 4.17 |
|  | Meridol | 1 | 7.08 | 5.35 |
|  |  | 2 | 7.07 | 5.05 |
|  |  | 3 | 7.05 | 5.18 |
| Donor 10 | PBS | 1 | 7.05 | 4.24 |
|  |  | 2 | 7.06 | 4.24 |
|  |  | 3 | 7.04 | 4.28 |
|  | Elmex | 1 | 7.05 | 6.14 |
|  |  | 2 | 7.04 | 6.31 |
|  |  | 3 | 6.99 | 6.23 |
|  | Listerine | 1 | 6.87 | 4.21 |
|  |  | 2 | 7.02 | 4.34 |
|  |  | 3 | 7 | 4.24 |
|  | Meridol | 1 | 7.03 | 6.19 |
|  |  | 2 | 7.06 | 6.52 |
|  |  | 3 | 7.04 | 6.44 |
| Donor 11 | PBS | 1 | 7.02 | 4.51 |
|  |  | 2 | 7.02 | 4.67 |
|  |  | 3 | 6.91 | 5.34 |
|  | Elmex | 1 | 7.03 | 6.57 |
|  |  | 2 | 7.03 | 6.69 |
|  |  | 3 | 7.02 | 6.43 |
|  | Listerine | 1 | 7 | 4.54 |
|  |  | 2 | 7.02 | 4.66 |
|  |  | 3 | 7.04 | 4.58 |
|  | Meridol | 1 | 6.94 | 6.57 |
|  |  | 2 | 7.03 | 6.36 |
|  |  | 3 | 7.04 | 6.36 |
| Donor 12 | PBS | 1 | 6.95 | 4.35 |
|  |  | 2 | 6.95 | 4.35 |
|  |  | 3 | 6.91 | 4.36 |
|  | Elmex | 1 | 6.96 | 4.63 |
|  |  | 2 | 6.95 | 4.64 |
|  |  | 3 | 6.95 | 4.78 |
|  | Listerine | 1 | 6.95 | 4.31 |
|  |  | 2 | 6.95 | 4.32 |
|  |  | 3 | 6.99 | 4.32 |
|  | Meridol | 1 | 6.95 | 4.57 |
|  |  | 2 | 6.99 | 4.59 |
|  |  | 3 | 6.95 | 4.62 |
| Donor 13 | PBS | 1 | 6.94 | 4.47 |
|  |  | 2 | 6.95 | 4.44 |
|  |  | 3 | 6.95 | 4.45 |
|  | Elmex | 1 | 6.95 | 4.88 |
|  |  | 2 | 6.99 | 4.79 |
|  |  | 3 | 6.95 | 4.87 |
|  | Listerine | 1 | 6.96 | 4.41 |
|  |  | 2 | 6.99 | 4.43 |
|  |  | 3 | 6.99 | 4.45 |
|  | Meridol | 1 | 6.99 | 5 |
|  |  | 2 | 6.95 | 4.95 |
|  |  | 3 | 6.99 | 5.06 |

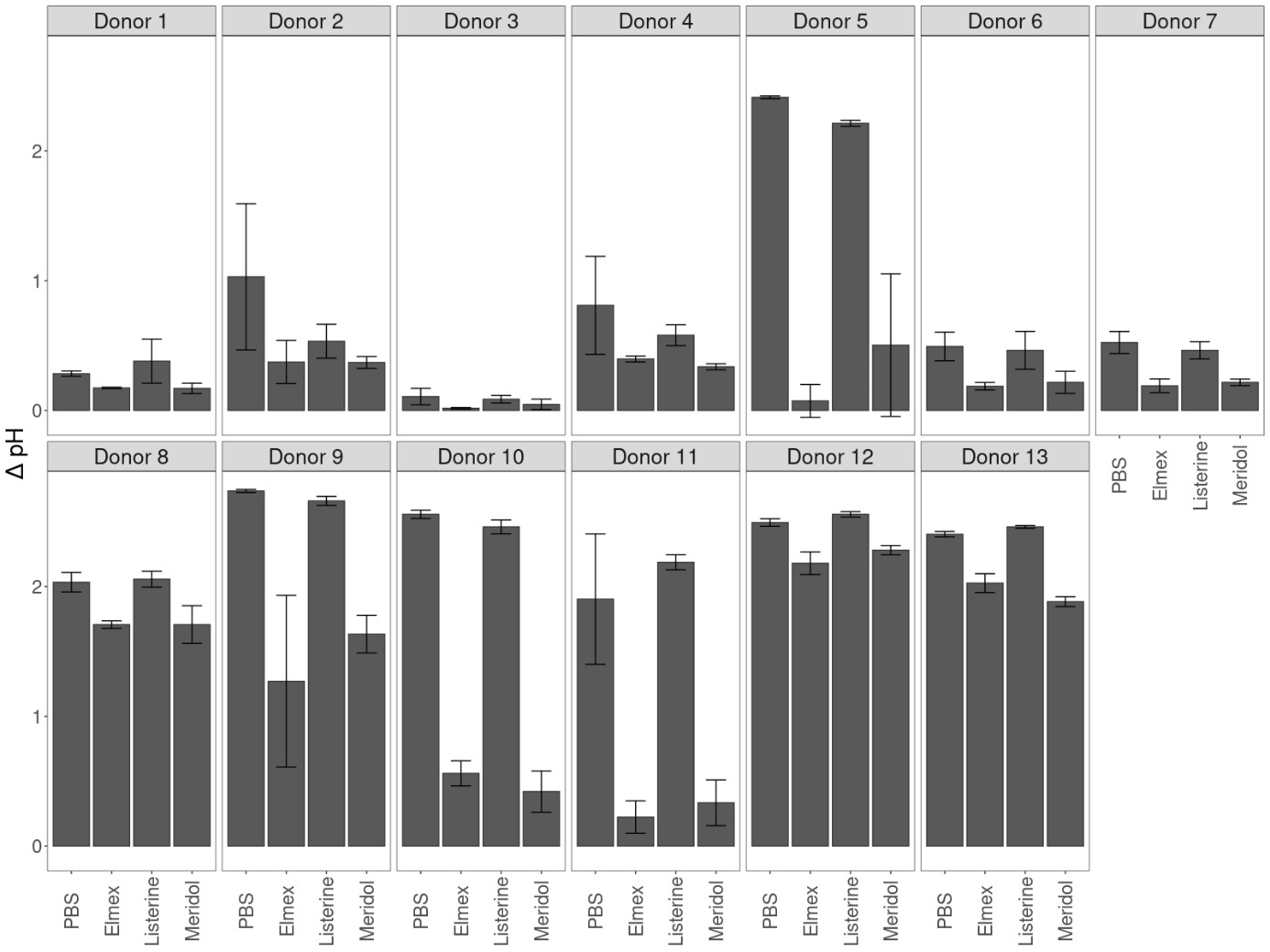

Figure S1. Drop in pH over microaerophilic incubation of mouthwash-treated saliva samples in minimal medium with sucrose as carbon source. Error bars indicate the standard deviation.

**Supplementary 4. Information on FACS**

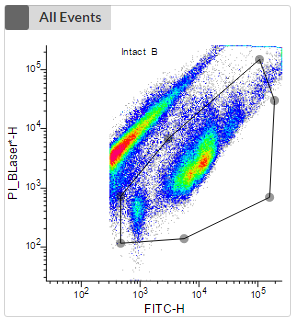

Figure S2. Gating strategy for FACS. Control samples (untreated saliva, filter-sterilized saliva, heat-killed saliva, sterile PBS, sterile medium) were used to determine the intact cell population. The intact cell population was gated based on green fluorescence (PMT1, ‘FITC-H’) and red fluorescence (PMT2, ‘PI_BLaser-H’).

Table S4. Information on the number of events sorted for the intact cell gate for each sample. Samples were sorted immediately after treatment. The target count was set at 250000 events.

| **Donor** | **Treatment** | **Replicate** | **Number of sorted events** |
| --- | --- | --- | --- |
| Donor 1 | Elmex | 3 | 7772 |
|  | Listerine | 3 | 250000 |
|  | Meridol | 3 | 8475 |
|  | PBS | 3 | 250000 |
| Donor 2 | Elmex | 1 | 250000 |
|  |  | 2 | 250000 |
|  |  | 3 | 250000 |
|  | Listerine | 1 | 250000 |
|  |  | 2 | 250000 |
|  |  | 3 | 250000 |
|  | Meridol | 1 | 250000 |
|  |  | 2 | 250000 |
|  |  | 3 | 250000 |
|  | PBS | 1 | 250000 |
|  |  | 2 | 250000 |
|  |  | 3 | 250000 |
| Donor 4 | Elmex | 2 | 130064 |
|  | Listerine | 2 | 250000 |
|  | Meridol | 2 | 250000 |
|  | PBS | 2 | 250000 |
| Donor 5 | Elmex | 1 | 50713 |
|  | Listerine | 1 | 250000 |
|  | Meridol | 1 | 76971 |
|  | PBS | 1 | 250000 |
| Donor 6 | Elmex | 1 | 34047 |
|  | Listerine | 1 | 250000 |
|  | Meridol | 1 | 118517 |
|  | PBS | 1 | 250000 |
| Donor 7 | Elmex | 3 | 77188 |
|  | Listerine | 3 | 250000 |
|  | Meridol | 3 | 94426 |
|  | PBS | 3 | 250000 |
| Donor 8 | Elmex | 1 | 250000 |
|  |  | 2 | 250000 |
|  |  | 3 | 250000 |
|  | Listerine | 1 | 250000 |
|  |  | 2 | 250000 |
|  |  | 3 | 250000 |
|  | Meridol | 1 | 250000 |
|  |  | 2 | 250000 |
|  |  | 3 | 250000 |
|  | PBS | 1 | 250000 |
|  |  | 2 | 250000 |
|  |  | 3 | 250000 |
| Donor 9 | Elmex | 1 | 250000 |
|  | Listerine | 1 | 250000 |
|  | Meridol | 1 | 250000 |
|  | PBS | 1 | 250000 |
| Donor 10 | Elmex | 1 | 176757 |
|  | Listerine | 1 | 250000 |
|  | Meridol | 1 | 217266 |
|  | PBS | 1 | 250000 |
| Donor 11 | Elmex | 1 | 214176 |
|  |  | 2 | 221519 |
|  |  | 3 | 238850 |
|  | Listerine | 1 | 250000 |
|  |  | 2 | 250000 |
|  |  | 3 | 250000 |
|  | Meridol | 1 | 215758 |
|  |  | 2 | 250000 |
|  |  | 3 | 192032 |
|  | PBS | 1 | 250000 |
|  |  | 2 | 250000 |
|  |  | 3 | 250000 |
| Donor 12 | Elmex | 1 | 250000 |
|  |  | 2 | 250000 |
|  |  | 3 | 250000 |
|  | Listerine | 1 | 250000 |
|  |  | 2 | 250000 |
|  |  | 3 | 250000 |
|  | Meridol | 1 | 250000 |
|  |  | 2 | 250000 |
|  |  | 3 | 250000 |
|  | PBS | 1 | 250000 |
|  |  | 2 | 250000 |
|  |  | 3 | 250000 |
| Donor 13 | Elmex | 2 | 187713 |
|  | Listerine | 2 | 250000 |
|  | Meridol | 2 | 223581 |
|  | PBS | 2 | 250000 |

**Supplementary 5. Overview of sequenced samples**

Table S5. Overview of samples that were sequenced using 16S rRNA gene amplicon sequencing.

| **Timepoint** | **Donor** | **Treatment** | **Population** | **Replicate** | **Raw read count** | **Read count after chimera removal** | **ASVs** |
| --- | --- | --- | --- | --- | --- | --- | --- |
| 0h | Donor 1 | Elmex | Intact | 3 | 1067 | 770 | 27 |
|  |  | Listerine | Intact | 3 | 12373 | 8032 | 85 |
|  |  | Meridol | Intact | 3 | 1608 | 1042 | 35 |
|  |  | PBS | Intact | 3 | 21471 | 14152 | 212 |
|  |  | Raw | Whole | 1 | 72028 | 47457 | 231 |
|  | Donor 2 | Elmex | Intact | 1 | 99345 | 69379 | 254 |
|  |  |  |  | 2 | 89844 | 67580 | 272 |
|  |  |  |  | 3 | 4766 | 3198 | 74 |
|  |  | Listerine | Intact | 1 | 22348 | 16060 | 149 |
|  |  |  |  | 2 | 68012 | 49017 | 208 |
|  |  |  |  | 3 | 21655 | 13259 | 92 |
|  |  | Meridol | Intact | 1 | 14185 | 9687 | 110 |
|  |  |  |  | 2 | 17445 | 11475 | 89 |
|  |  |  |  | 3 | 3950 | 2521 | 30 |
|  |  | PBS | Intact | 1 | 20777 | 13358 | 125 |
|  |  |  |  | 2 | 12019 | 8379 | 133 |
|  |  |  |  | 3 | 111645 | 78972 | 255 |
|  |  | Raw | Whole | 1 | 96984 | 60174 | 183 |
|  | Donor 3 | Raw | Whole | 1 | 66566 | 43151 | 233 |
|  | Donor 4 | Elmex | Intact | 2 | 81532 | 56264 | 302 |
|  |  | Listerine | Intact | 2 | 22763 | 16157 | 195 |
|  |  | Meridol | Intact | 2 | 40300 | 27539 | 218 |
|  |  | PBS | Intact | 2 | 31647 | 19303 | 217 |
|  |  | Raw | Whole | 1 | 50062 | 31333 | 162 |
|  | Donor 5 | Elmex | Intact | 1 | 389 | 41 | 4 |
|  |  | Listerine | Intact | 1 | 34041 | 20608 | 103 |
|  |  | Meridol | Intact | 1 | 2207 | 1510 | 38 |
|  |  | PBS | Intact | 1 | 4572 | 2924 | 58 |
|  |  | Raw | Whole | 1 | 72648 | 46001 | 171 |
|  | Donor 6 | Elmex | Intact | 1 | 406 | 219 | 22 |
|  |  | Listerine | Intact | 1 | 90721 | 59821 | 261 |
|  |  | Meridol | Intact | 1 | 6293 | 3693 | 62 |
|  |  | PBS | Intact | 1 | 5233 | 3404 | 97 |
|  |  | Raw | Whole | 1 | 14043 | 9157 | 105 |
|  | Donor 7 | Elmex | Intact | 3 | 328 | 142 | 14 |
|  |  | Listerine | Intact | 3 | 62262 | 41184 | 246 |
|  |  | Meridol | Intact | 3 | 10724 | 6772 | 113 |
|  |  | PBS | Intact | 3 | 1694 | 909 | 33 |
|  |  | Raw | Whole | 1 | 67450 | 45385 | 116 |
|  | Donor 8 | Elmex | Intact | 1 | 4376 | 2844 | 60 |
|  |  |  |  | 2 | 39003 | 26559 | 199 |
|  |  |  |  | 3 | 35894 | 25078 | 179 |
|  |  | Listerine | Intact | 1 | 88912 | 60208 | 265 |
|  |  |  |  | 2 | 56900 | 42296 | 267 |
|  |  |  |  | 3 | 14973 | 9542 | 107 |
|  |  | Meridol | Intact | 1 | 40112 | 26873 | 186 |
|  |  |  |  | 2 | 24476 | 13020 | 125 |
|  |  |  |  | 3 | 14390 | 8927 | 111 |
|  |  | PBS | Intact | 1 | 112407 | 66608 | 247 |
|  |  |  |  | 2 | 56816 | 36871 | 234 |
|  |  |  |  | 3 | 15282 | 9616 | 109 |
|  |  | Raw | Whole | 1 | 51746 | 34478 | 134 |
|  | Donor 9 | Elmex | Intact | 1 | 13512 | 9430 | 117 |
|  |  | Listerine | Intact | 1 | 95323 | 77218 | 303 |
|  |  | Meridol | Intact | 1 | 36638 | 25413 | 197 |
|  |  | PBS | Intact | 1 | 21427 | 14289 | 155 |
|  |  | Raw | Whole | 1 | 92260 | 55886 | 168 |
|  | Donor 10 | Elmex | Intact | 1 | 56576 | 36152 | 124 |
|  |  | Listerine | Intact | 1 | 21777 | 14356 | 97 |
|  |  | Meridol | Intact | 1 | 20343 | 13337 | 96 |
|  |  | PBS | Intact | 1 | 67327 | 40482 | 123 |
|  |  | Raw | Whole | 1 | 52822 | 35590 | 118 |
|  | Donor 11 | Elmex | Intact | 1 | 36967 | 24182 | 245 |
|  |  |  |  | 2 | 13154 | 7558 | 108 |
|  |  |  |  | 3 | 1959 | 1154 | 33 |
|  |  | Listerine | Intact | 1 | 30135 | 16224 | 137 |
|  |  |  |  | 2 | 19952 | 13380 | 173 |
|  |  |  |  | 3 | 7086 | 4187 | 70 |
|  |  | Meridol | Intact | 1 | 20242 | 14908 | 186 |
|  |  |  |  | 2 | 5955 | 3647 | 86 |
|  |  |  |  | 3 | 14457 | 7456 | 104 |
|  |  | PBS | Intact | 1 | 34285 | 24636 | 258 |
|  |  |  |  | 2 | 10342 | 7025 | 140 |
|  |  |  |  | 3 | 67035 | 35698 | 253 |
|  |  | Raw | Whole | 1 | 61053 | 37095 | 170 |
|  | Donor 12 | Elmex | Intact | 1 | 19352 | 12810 | 123 |
|  |  |  |  | 2 | 43753 | 31576 | 213 |
|  |  |  |  | 3 | 53573 | 36420 | 180 |
|  |  | Listerine | Intact | 1 | 11347 | 7700 | 96 |
|  |  |  |  | 2 | 15525 | 10994 | 130 |
|  |  |  |  | 3 | 20913 | 12911 | 104 |
|  |  | Meridol | Intact | 1 | 40419 | 28856 | 210 |
|  |  |  |  | 2 | 45475 | 30703 | 213 |
|  |  |  |  | 3 | 27669 | 19070 | 177 |
|  |  | PBS | Intact | 1 | 51222 | 34789 | 199 |
|  |  |  |  | 2 | 162473 | 127341 | 318 |
|  |  |  |  | 3 | 25200 | 17642 | 176 |
|  |  | Raw | Whole | 1 | 30502 | 18803 | 127 |
|  | Donor 13 | Elmex | Intact | 2 | 108228 | 81345 | 354 |
|  |  | Listerine | Intact | 2 | 29813 | 18434 | 145 |
|  |  | Meridol | Intact | 2 | 26572 | 18908 | 199 |
|  |  | PBS | Intact | 2 | 5694 | 3687 | 67 |
|  |  | Raw | Whole | 1 | 85260 | 55040 | 227 |
| 19h | Donor 1 | Elmex | Whole | 1 | 52040 | 43134 | 189 |
|  |  | Listerine | Whole | 1 | 24793 | 20038 | 101 |
|  |  | Meridol | Whole | 1 | 6982 | 5492 | 51 |
|  |  | PBS | Whole | 1 | 60865 | 50159 | 130 |
|  | Donor 2 | Elmex | Whole | 1 | 94045 | 78432 | 35 |
|  |  |  |  | 2 | 67138 | 50433 | 157 |
|  |  |  |  | 3 | 84093 | 68787 | 115 |
|  |  | Listerine | Whole | 1 | 95912 | 75998 | 92 |
|  |  |  |  | 2 | 176704 | 140001 | 99 |
|  |  |  |  | 3 | 50919 | 40488 | 71 |
|  |  | Meridol | Whole | 1 | 54598 | 43879 | 97 |
|  |  |  |  | 2 | 88089 | 66961 | 89 |
|  |  |  |  | 3 | 72217 | 57822 | 21 |
|  |  | PBS | Whole | 1 | 148137 | 117771 | 65 |
|  |  |  |  | 2 | 132124 | 106614 | 84 |
|  |  |  |  | 3 | 94470 | 76931 | 97 |
|  | Donor 3 | Elmex | Whole | 1 | 30476 | 25171 | 97 |
|  |  |  |  | 2 | 17432 | 14448 | 87 |
|  |  |  |  | 3 | 19378 | 16322 | 84 |
|  |  | Listerine | Whole | 1 | 3978 | 3082 | 43 |
|  |  |  |  | 2 | 5141 | 4143 | 50 |
|  |  |  |  | 3 | 23004 | 18524 | 96 |
|  |  | Meridol | Whole | 1 | 9720 | 8173 | 65 |
|  |  |  |  | 2 | 2449 | 1931 | 37 |
|  |  |  |  | 3 | 1722 | 1385 | 30 |
|  |  | PBS | Whole | 1 | 22350 | 18341 | 125 |
|  |  |  |  | 2 | 96493 | 74930 | 141 |
|  |  |  |  | 3 | 8402 | 6506 | 92 |
|  | Donor 4 | Elmex | Whole | 1 | 58073 | 43818 | 158 |
|  |  | Listerine | Whole | 1 | 13272 | 10686 | 44 |
|  |  | Meridol | Whole | 1 | 27089 | 22228 | 116 |
|  |  | PBS | Whole | 1 | 38211 | 30163 | 88 |
|  | Donor 5 | Elmex | Whole | 1 | 22198 | 18037 | 37 |
|  |  | Listerine | Whole | 1 | 59996 | 47770 | 24 |
|  |  | Meridol | Whole | 1 | 45295 | 37562 | 53 |
|  |  | PBS | Whole | 1 | 61698 | 50816 | 36 |
|  | Donor 6 | Elmex | Whole | 1 | 19165 | 15835 | 81 |
|  |  | Listerine | Whole | 1 | 61356 | 41342 | 89 |
|  |  | Meridol | Whole | 1 | 37284 | 28081 | 47 |
|  |  | PBS | Whole | 1 | 81073 | 53079 | 93 |
|  | Donor 7 | Elmex | Whole | 1 | 26594 | 21712 | 78 |
|  |  | Listerine | Whole | 1 | 19657 | 15542 | 30 |
|  |  | Meridol | Whole | 1 | 3354 | 2706 | 12 |
|  |  | PBS | Whole | 1 | 46292 | 37324 | 51 |
|  | Donor 8 | Elmex | Whole | 1 | 41951 | 32566 | 48 |
|  |  | Listerine | Whole | 1 | 52232 | 40375 | 50 |
|  |  | Meridol | Whole | 1 | 62120 | 45862 | 35 |
|  |  | PBS | Whole | 1 | 57990 | 42574 | 57 |
|  | Donor 9 | Elmex | Whole | 1 | 72698 | 56235 | 67 |
|  |  | Listerine | Whole | 1 | 48322 | 39737 | 24 |
|  |  | Meridol | Whole | 1 | 20769 | 16421 | 27 |
|  |  | PBS | Whole | 1 | 61219 | 48888 | 25 |
|  | Donor 10 | Elmex | Whole | 1 | 56262 | 46776 | 28 |
|  |  | Listerine | Whole | 1 | 69734 | 56148 | 18 |
|  |  | Meridol | Whole | 1 | 23419 | 19690 | 13 |
|  |  | PBS | Whole | 1 | 28814 | 23759 | 9 |
|  | Donor 11 | Elmex | Whole | 1 | 47152 | 37803 | 76 |
|  |  |  |  | 2 | 11366 | 9302 | 44 |
|  |  |  |  | 3 | 13816 | 11547 | 18 |
|  |  | Listerine | Whole | 1 | 27636 | 22197 | 10 |
|  |  |  |  | 2 | 17891 | 14661 | 10 |
|  |  |  |  | 3 | 36332 | 30268 | 18 |
|  |  | Meridol | Whole | 1 | 21268 | 17510 | 36 |
|  |  |  |  | 2 | 53653 | 43272 | 19 |
|  |  |  |  | 3 | 45056 | 36822 | 29 |
|  |  | PBS | Whole | 1 | 49245 | 40803 | 18 |
|  |  |  |  | 2 | 96698 | 77176 | 19 |
|  |  |  |  | 3 | 69198 | 55077 | 9 |
|  | Donor 12 | Elmex | Whole | 1 | 10514 | 8494 | 33 |
|  |  |  |  | 2 | 46559 | 36688 | 70 |
|  |  |  |  | 3 | 43577 | 35571 | 67 |
|  |  | Listerine | Whole | 1 | 43183 | 35252 | 50 |
|  |  |  |  | 2 | 35125 | 28880 | 39 |
|  |  |  |  | 3 | 24519 | 19782 | 35 |
|  |  | Meridol | Whole | 1 | 74417 | 57582 | 72 |
|  |  |  |  | 2 | 56039 | 43171 | 68 |
|  |  |  |  | 3 | 32632 | 26240 | 75 |
|  |  | PBS | Whole | 1 | 62642 | 50502 | 48 |
|  |  |  |  | 2 | 48713 | 40790 | 45 |
|  |  |  |  | 3 | 43331 | 35736 | 58 |
|  | Donor 13 | Elmex | Whole | 1 | 12661 | 10108 | 47 |
|  |  | Listerine | Whole | 1 | 36705 | 29841 | 53 |
|  |  | Meridol | Whole | 1 | 38859 | 30973 | 70 |
|  |  | PBS | Whole | 1 | 37592 | 31181 | 48 |
|  | Negative control | Medium batch 1 | Whole | 1 | 11 | 0 | 0 |
|  |  | Medium batch 2 | Whole | 1 | 14 | 0 | 0 |
|  |  | Medium batch 3 | Whole | 1 | 11 | 0 | 0 |
| *NA* | Negative control | Extraction blanc | Whole | 1 | 75 | 48 | 4 |
|  |  | Extraction blanc sorted | Whole | 1 | 922 | 594 | 10 |
|  |  |  |  | 2 | 455 | 317 | 12 |
|  |  | FACS Sheath | Whole | 1 | 1646 | 1012 | 9 |

**Supplementary 6. Gating strategy for flow cytometry analysis**

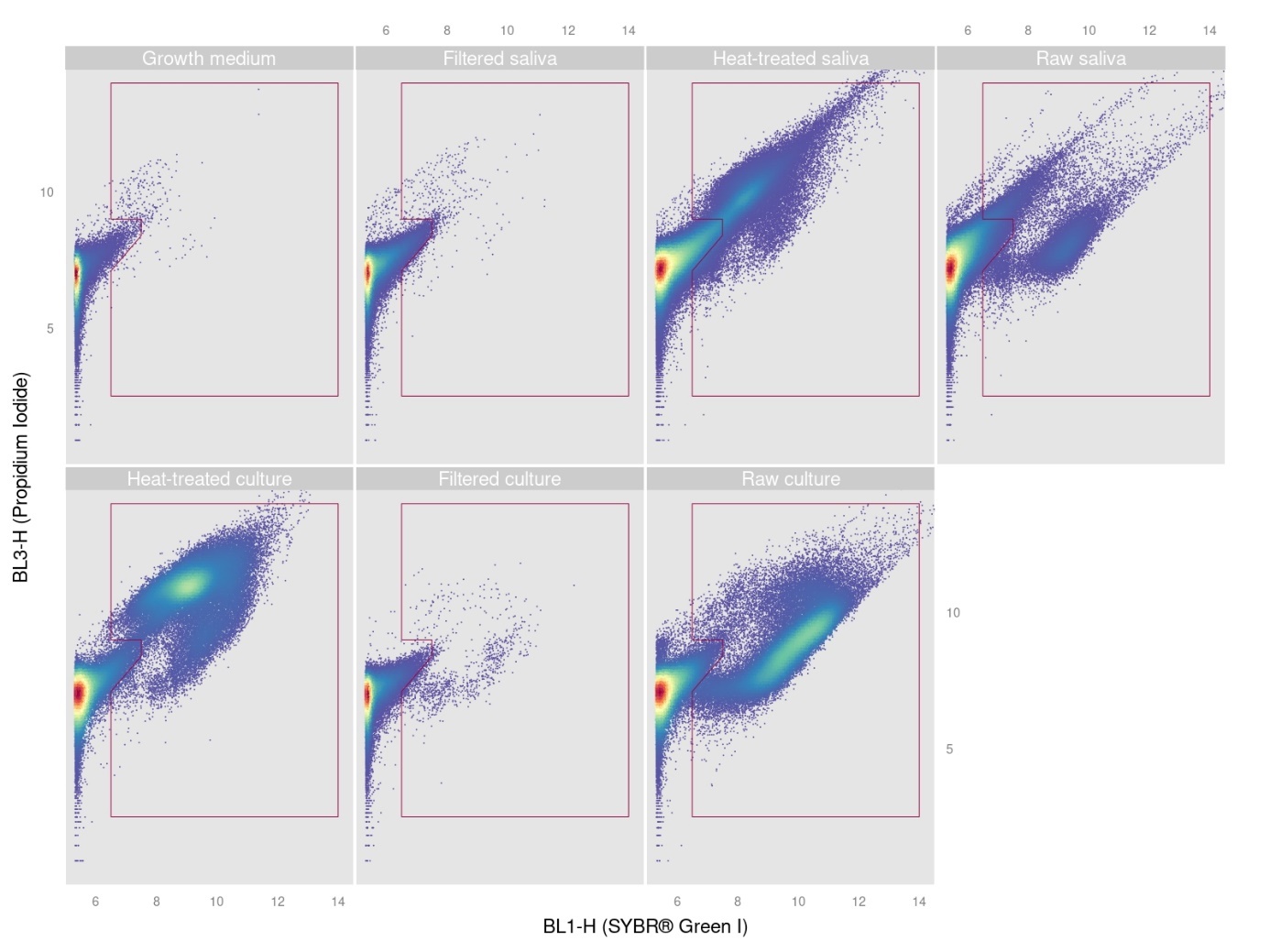

Figure S3. Gating strategy for gating total cells. Negative controls are represented by ‘Growth medium’, ‘Filtered saliva’ (pooled saliva sample from all donors that was filtered through a 0.2 µm filter), and ‘Filtered culture’ (pooled sample of 19h-incubated saliva that was filtered through a 0.2 µm filter). ‘Heat-treated saliva’ and ‘Heat-treated culture’ represent pooled samples that were microwaved, and ‘Raw saliva’ and ‘Raw culture’ represent untreated samples. The gate was drawn so that background was excluded and cells were included.

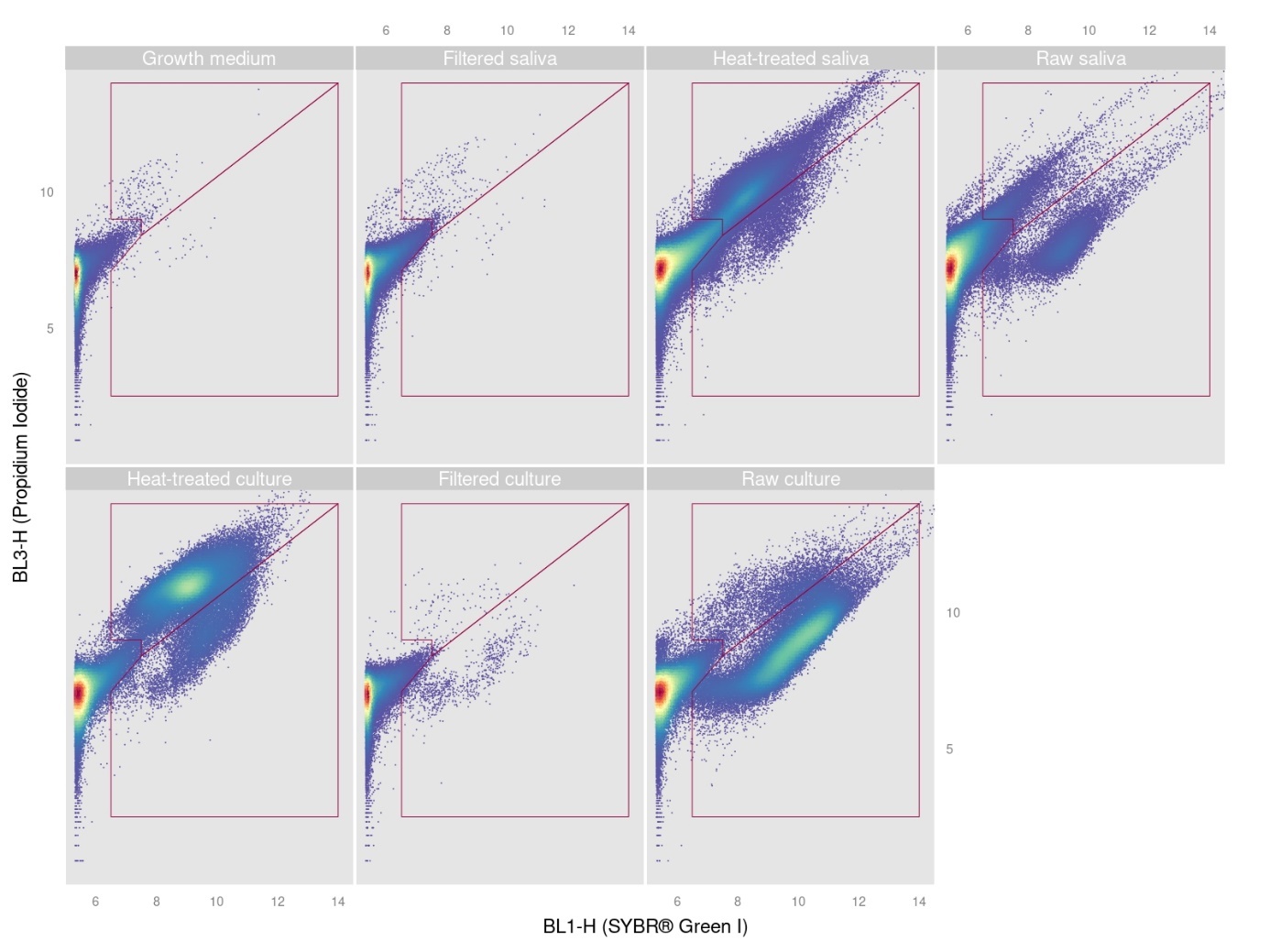

Figure S4. Gating strategy for gating intact (lower gate) and damaged (upper gate) cells. Negative controls are represented by ‘Growth medium’, ‘Filtered saliva’ (pooled saliva sample from all donors that was filtered through a 0.2 µm filter), and ‘Filtered culture’ (pooled sample of 19h-incubated saliva that was filtered through a 0.2 µm filter). ‘Heat-treated saliva’ and ‘Heat-treated culture’ represent pooled samples that were microwaved, and ‘Raw saliva’ and ‘Raw culture’ represent untreated samples.

**Supplementary 7. Construction of phenotypic fingerprints**

Fingerprints were constructed by applying a Gaussian mixture mask to the flow cytometric data. Donors and treatments were subsampled to an equal number of cells to avoid biased model training towards a specific donor or treatment. The number of clusters for the Gaussian mixture model (GMM) was decided by optimization of the Bayesian information criterion (BIC) using *PhenoGMM*. This was done for the intact cell population before incubation, the total cell population after incubation and the intact cell population of the PBS-treated samples before incubation separately. Figure S5 shows the BIC for different numbers of clusters included in the GMM. The number of clusters in the final model (i.e. the mask) was decided through determining for which number of clusters the BIC starts to converge to its maximum for most model types. The number of clusters was set to 100 for the final model for the intact cell population before incubation and to 60 for the final models for the total cell population after incubation and the intact cell population of the PBS-treated samples before incubation.

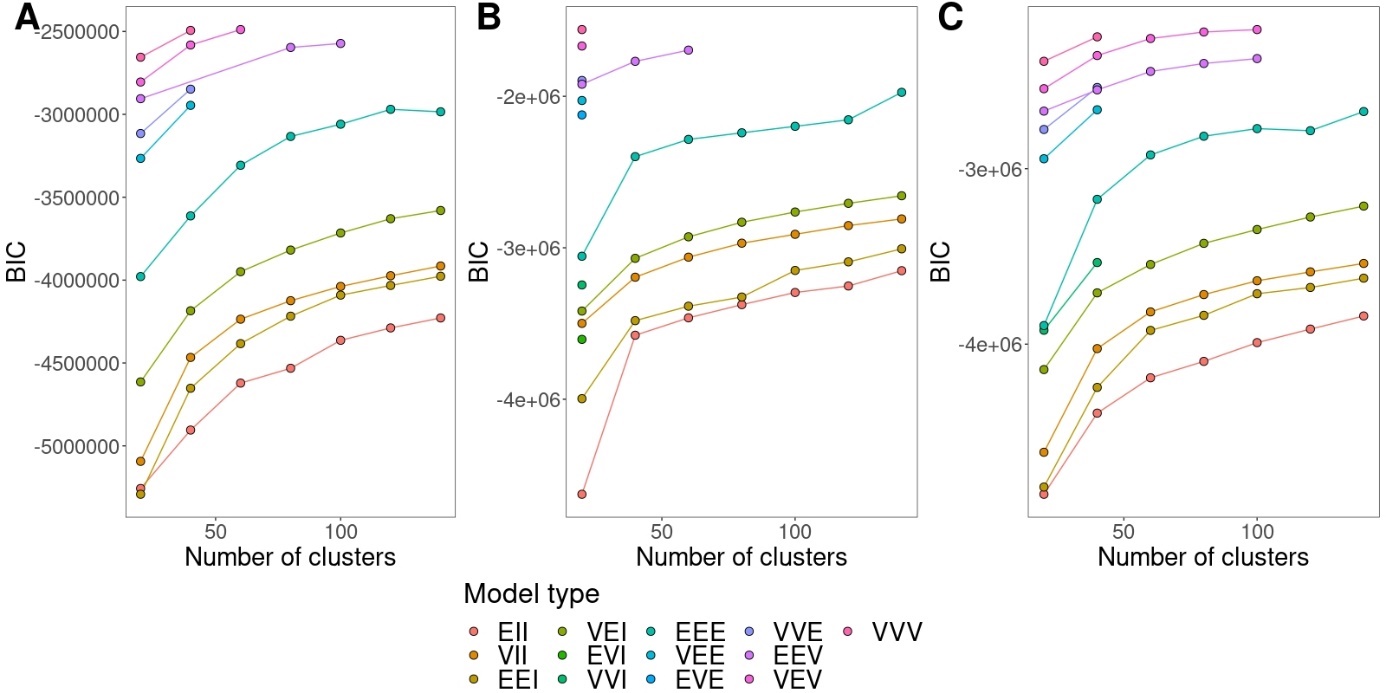

Figure S5. BIC values for different model types for the intact cell population before incubation (A), the total cell population after incubation (B) and the intact cell population of the PBS-treated samples before incubation (C).

**Supplementary 8. Rarefaction curves amplicon sequencing**

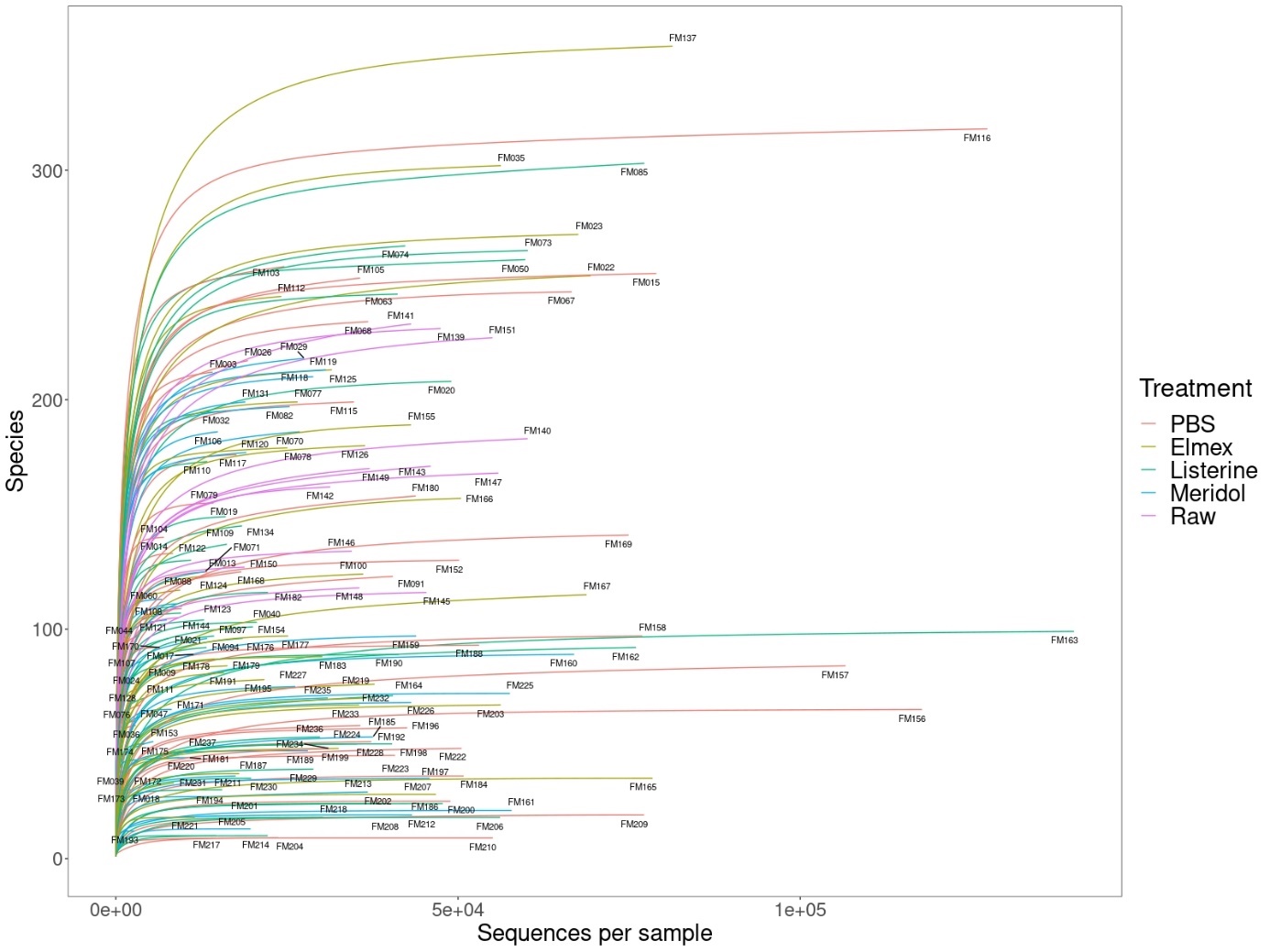

Figure S6. Rarefaction curves for all samples. Colors indicate the treatment.

**Supplementary 9. Cell concentrations determined by flow cytometry**

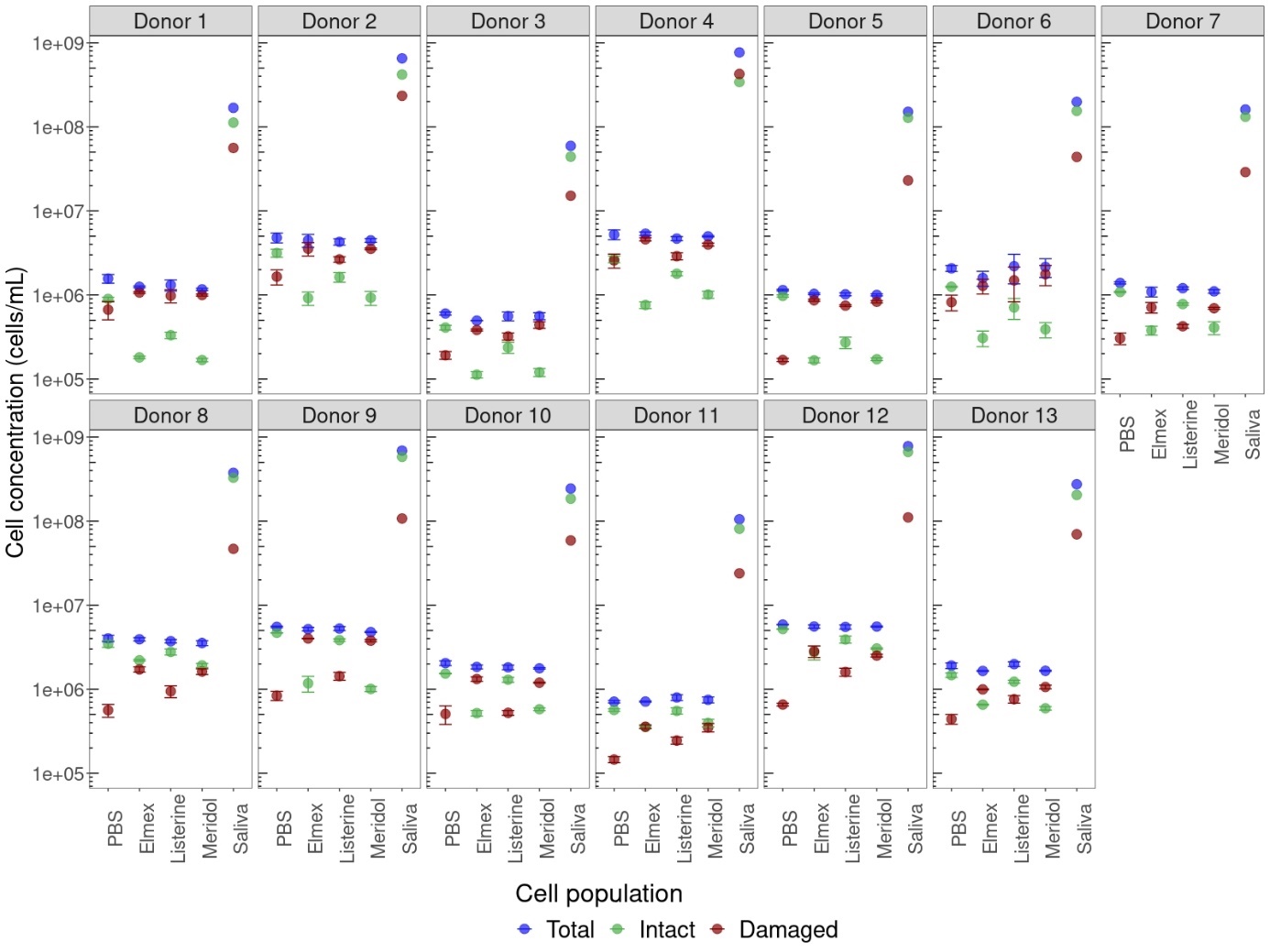

Figure S7. Absolute cell concentrations of saliva samples after treatment with different fluoride containing mouthwashes as determined by flow cytometry. ‘PBS’ indicates the control and ‘Saliva’ shows the cell concentrations of raw saliva. Error bars show the standard deviation.

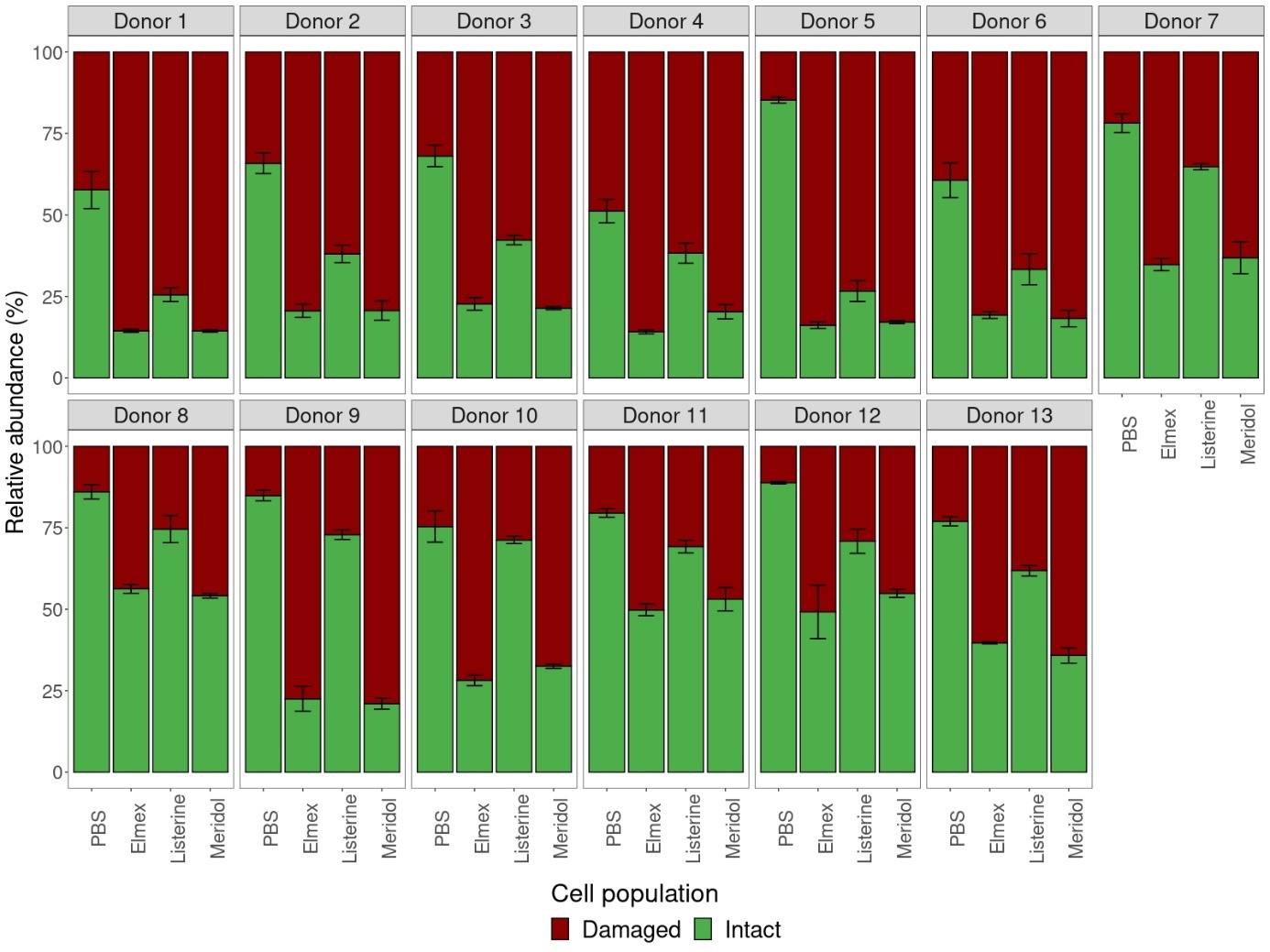

Figure S8. Relative abundance of intact and damaged cell populations of saliva samples after treatment with different fluoride containing mouthwashes as determined by flow cytometry. ‘PBS’ indicates the control. Error bars show the standard deviation.

**Supplementary 10. Quantitative community composition after mouthwash treatment**

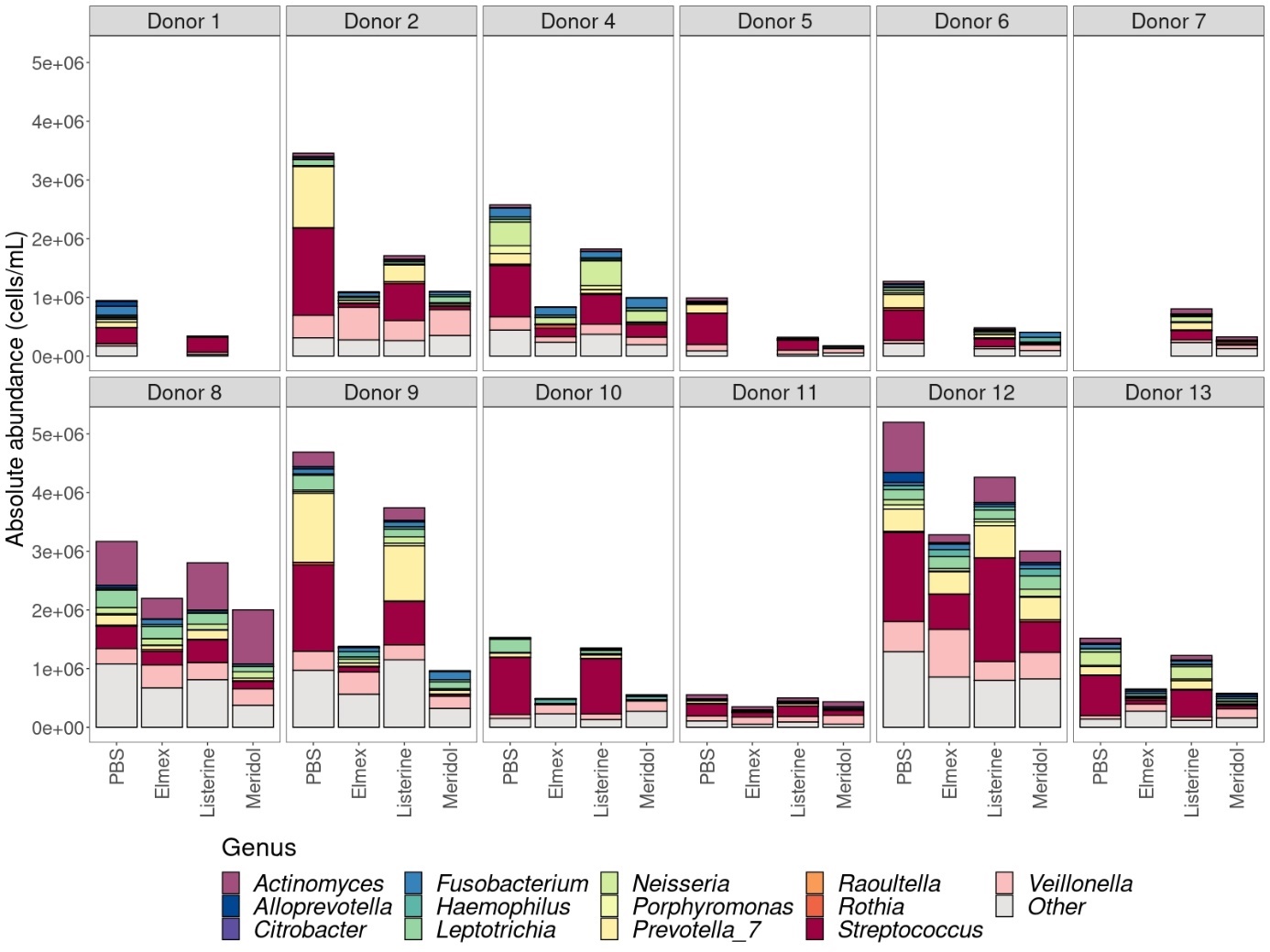

Figure S9. Quantitative community composition of the 13 most abundant bacterial genera in the intact cell population after mouthwash treatment. Lower abundant genera were grouped into ‘Other’. The missing samples are samples that were omitted from analysis due to a low number of read counts for these samples.

**Supplementary 11. Alpha diversity after mouthwash treatment**

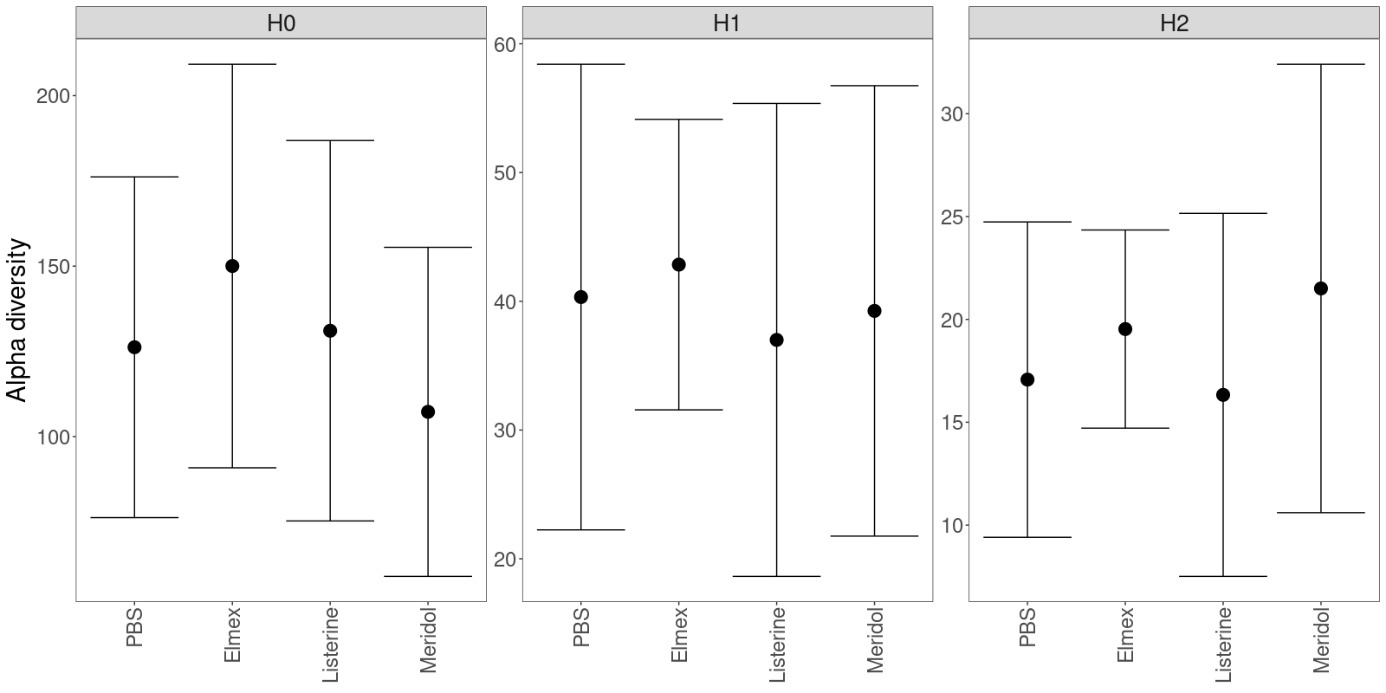

Figure S10. Mean α-diversity indices over all donors after mouthwash treatment. No significant differences between the different treatments were found for either species richness (H0), Shannon diversity (H1) and inverse Simpson diversity (H2). Statistical differences were assessed using mixed-effect models with Benjamini-Hochberg correction for multiple testing. Error bars indicate the standard deviation.

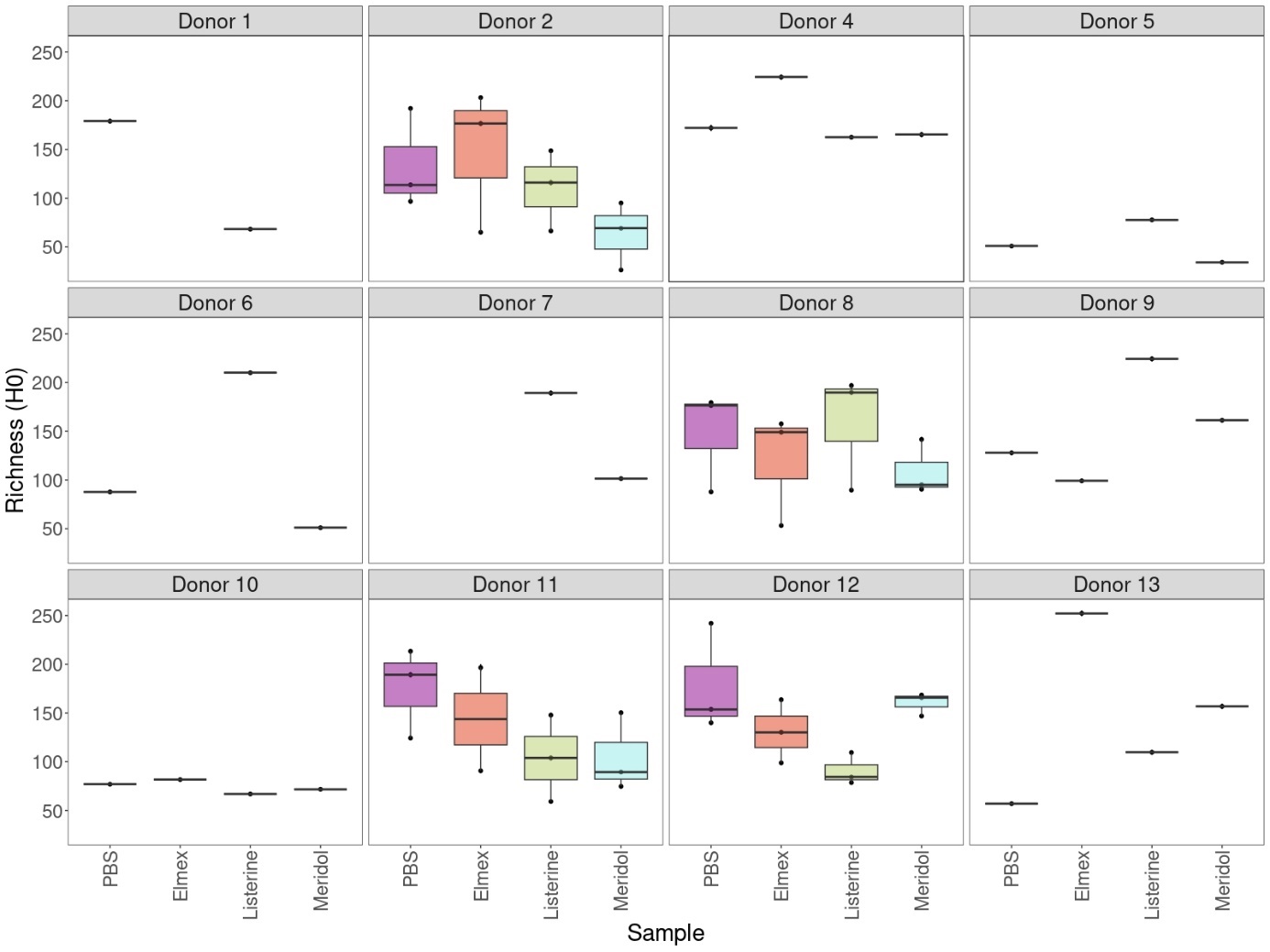

Figure S11. Species richness (H0) at extrapolated sample coverage for the intact cell population after mouthwash treatment for the different donors. The missing samples are samples that were omitted from analysis due to a low number of read counts for these samples.

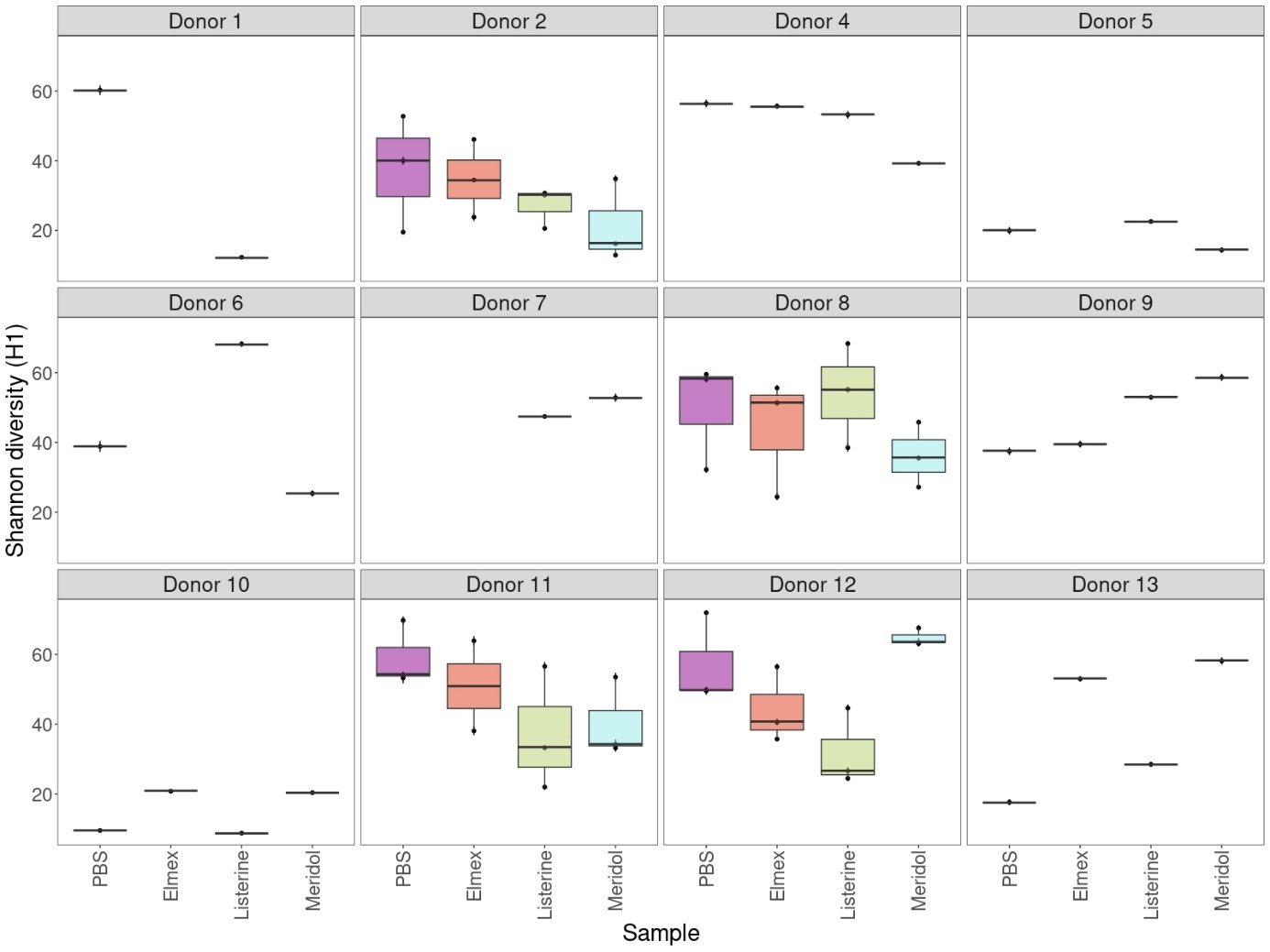

Figure S12. Shannon diversity (H1) at extrapolated sample coverage for the intact cell population after mouthwash treatment for the different donors. The missing samples are samples that were omitted from analysis due to a low number of read counts for these samples.

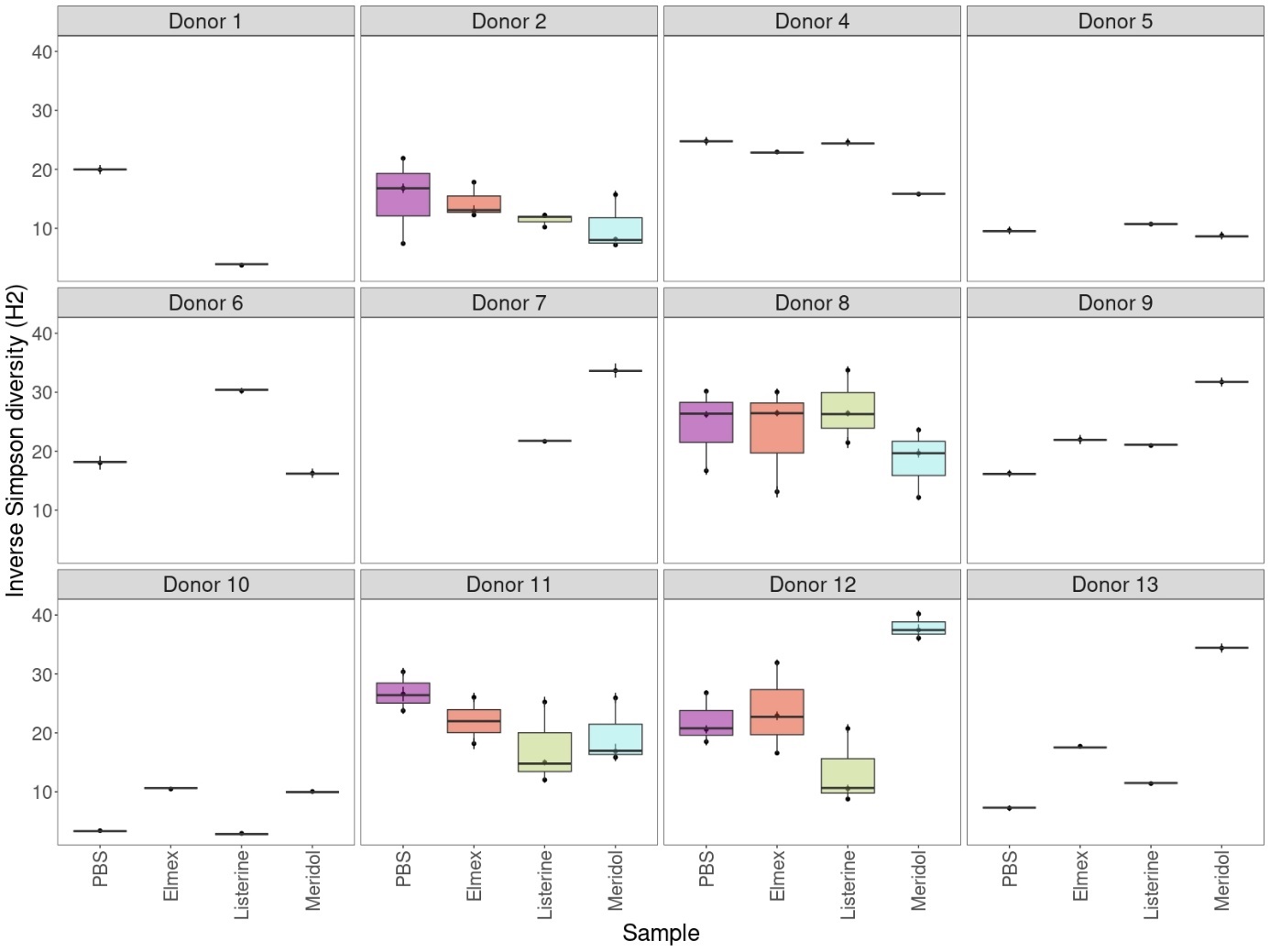

Figure S13. Inverse Simpson diversity (H2) at extrapolated sample coverage for the intact cell population after mouthwash treatment for the different donors. The missing samples are samples that were omitted from analysis due to a low number of read counts for these samples.

**Supplementary 12. PCoA flow cytometric fingerprints between treatments**

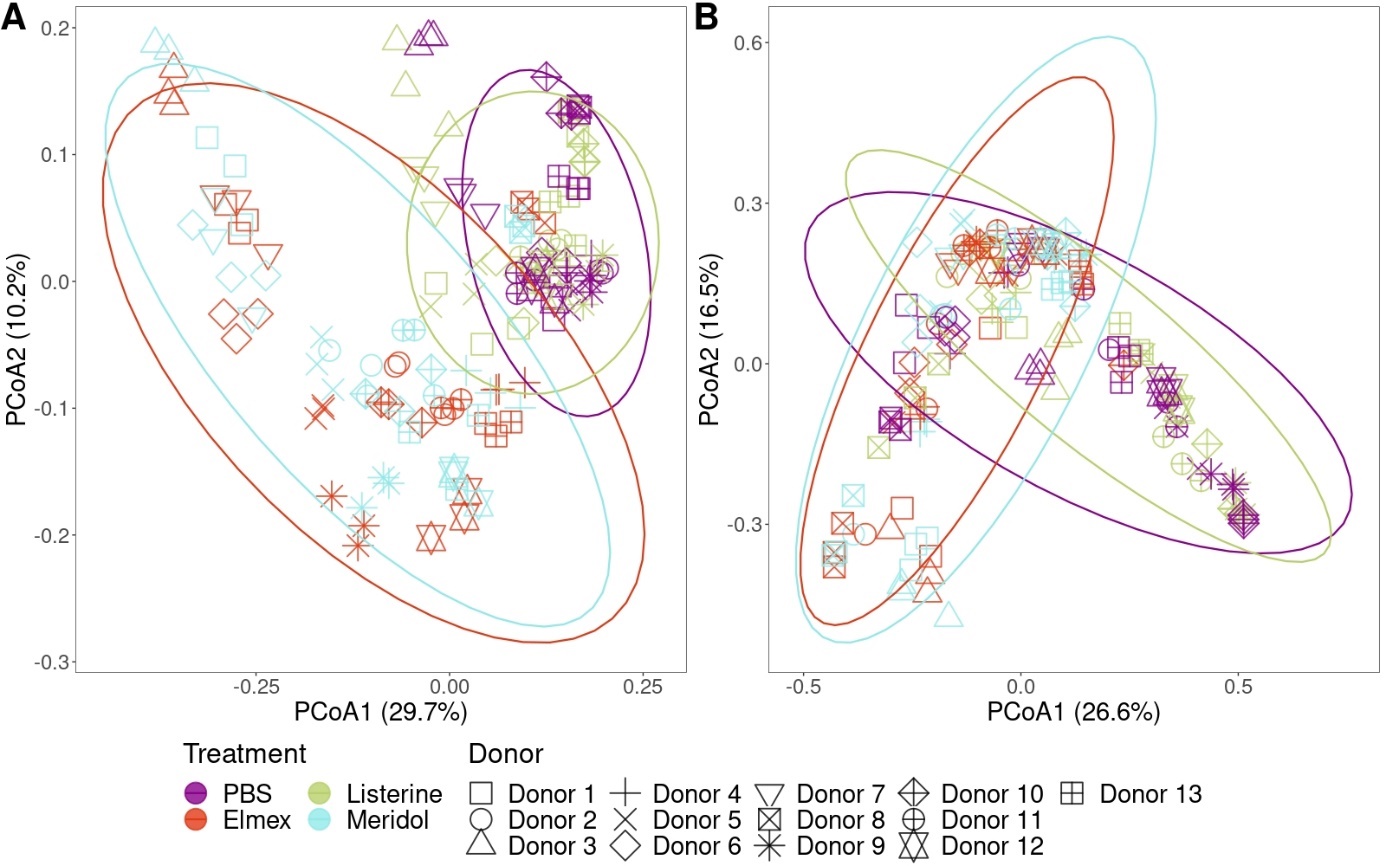

Figure S14. PCoA based on Bray-Curtis dissimilarity of the flow cytometric fingerprint of the intact cell population after mouthwash treatment (A) and of the total cell population after incubation (B). Ellipses were drawn at the 95% confidence level. For the intact cell population after mouthwash treatment, statistical differences were found between PBS and Elmex (ANOSIM; *R* = 0.51, *P* = 0.0001), PBS and Listerine (ANOSIM; *R* = 0.13, *P* = 0.0001), PBS and Meridol (ANOSIM; *R* = 0.56, *P* = 0.0001), Listerine and Elmex (ANOSIM; *R* = 0.4, *P* = 0.0001), and Listerine and Meridol (ANOSIM; *R* = 0.44, *P* = 0.0001). For the total cell population after incubation, statistical differences were found between PBS and Elmex (ANOSIM; *R* = 0.17, *P* = 0.0001), PBS and Meridol (ANOSIM; *R* = 0.15, *P* = 0.0001), Listerine and Elmex (ANOSIM; *R* = 0.18, *P* = 0.0001), and Listerine and Meridol (ANOSIM; *R* = 0.16, *P* = 0.0001).

**Supplementary 13. db-RDA of proportional and quantitative community composition**

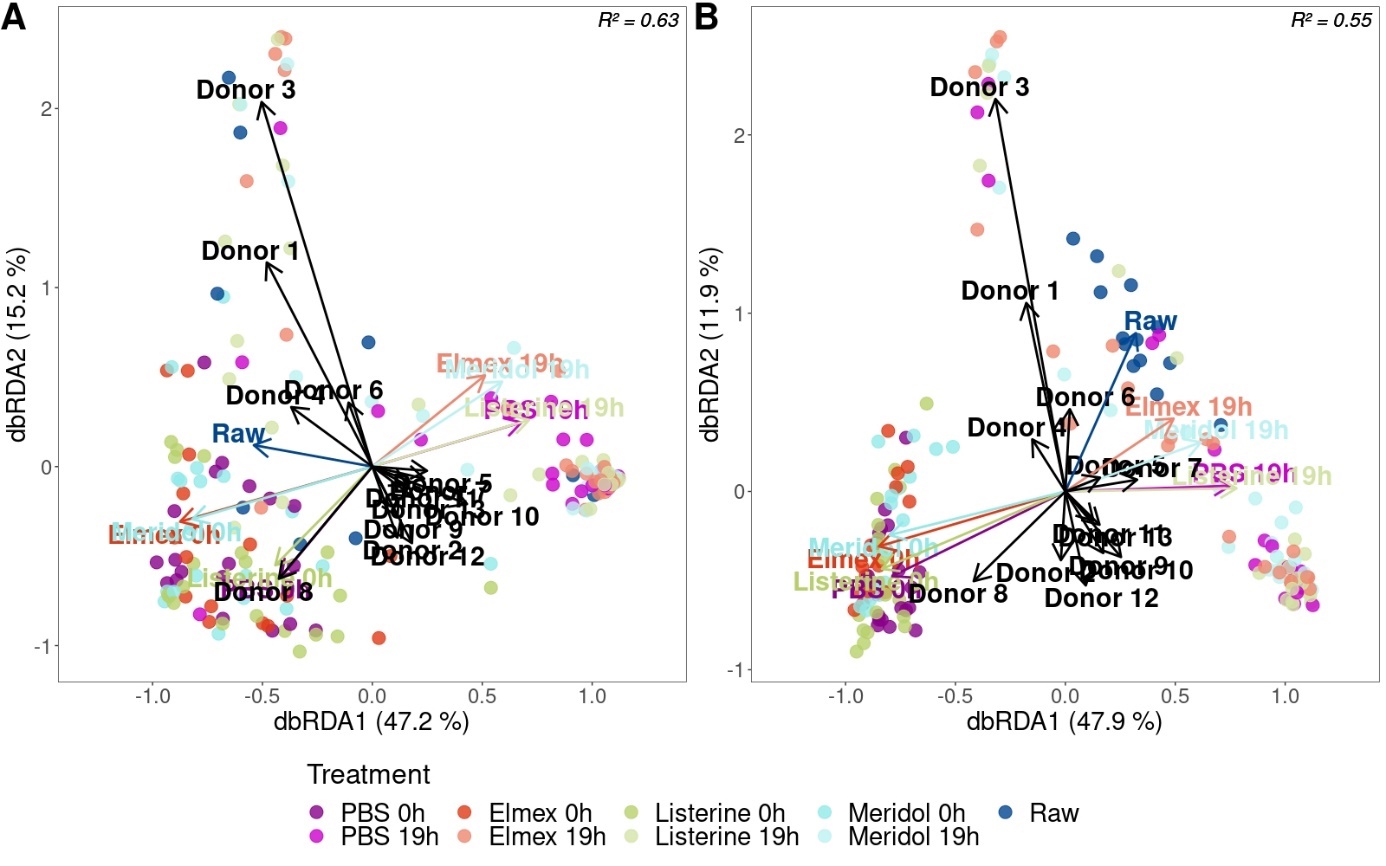

Figure S15. db-RDA triplots based on Bray-Curtis dissimilarity for the proportional (A) and quantitative (B) community composition (ASV level). Centroid factor levels are depicted by the arrows. Adjusted R² values in the top right corner of each plot show the model fit. Both ‘Treatment’ and ‘Donor’ contributed significantly to the variation in the model for the proportional (Treatment: *F* = 19.38, *P* = 0.001; Donor: *F* = 12.49, *P* = 0.001) and quantitative (Treatment: *F* = 16.50, *P* = 0.001; Donor: *F* = 7.99, *P* = 0.001) community data.

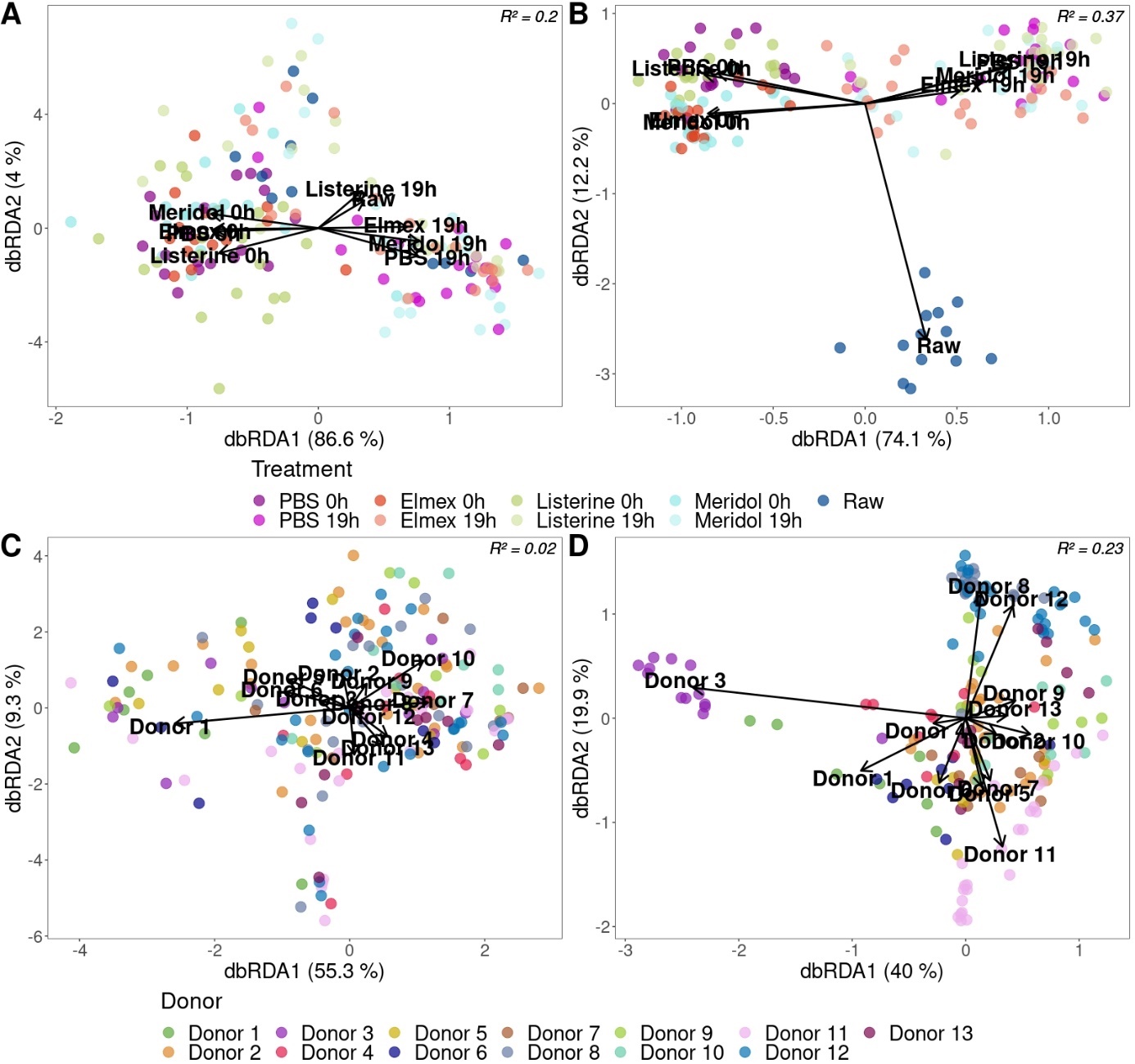

Figure S16. Partial db-RDA triplots based on Bray-Curtis dissimilarity of the proportional (A, C) and quantitative (B, D) community composition (ASV level). Centroid factor levels are depicted by the arrows. Adjusted R² values shown in the top right corner depict the model fit.

**Supplementary 14. Differential abundance analysis after mouthwash treatment**

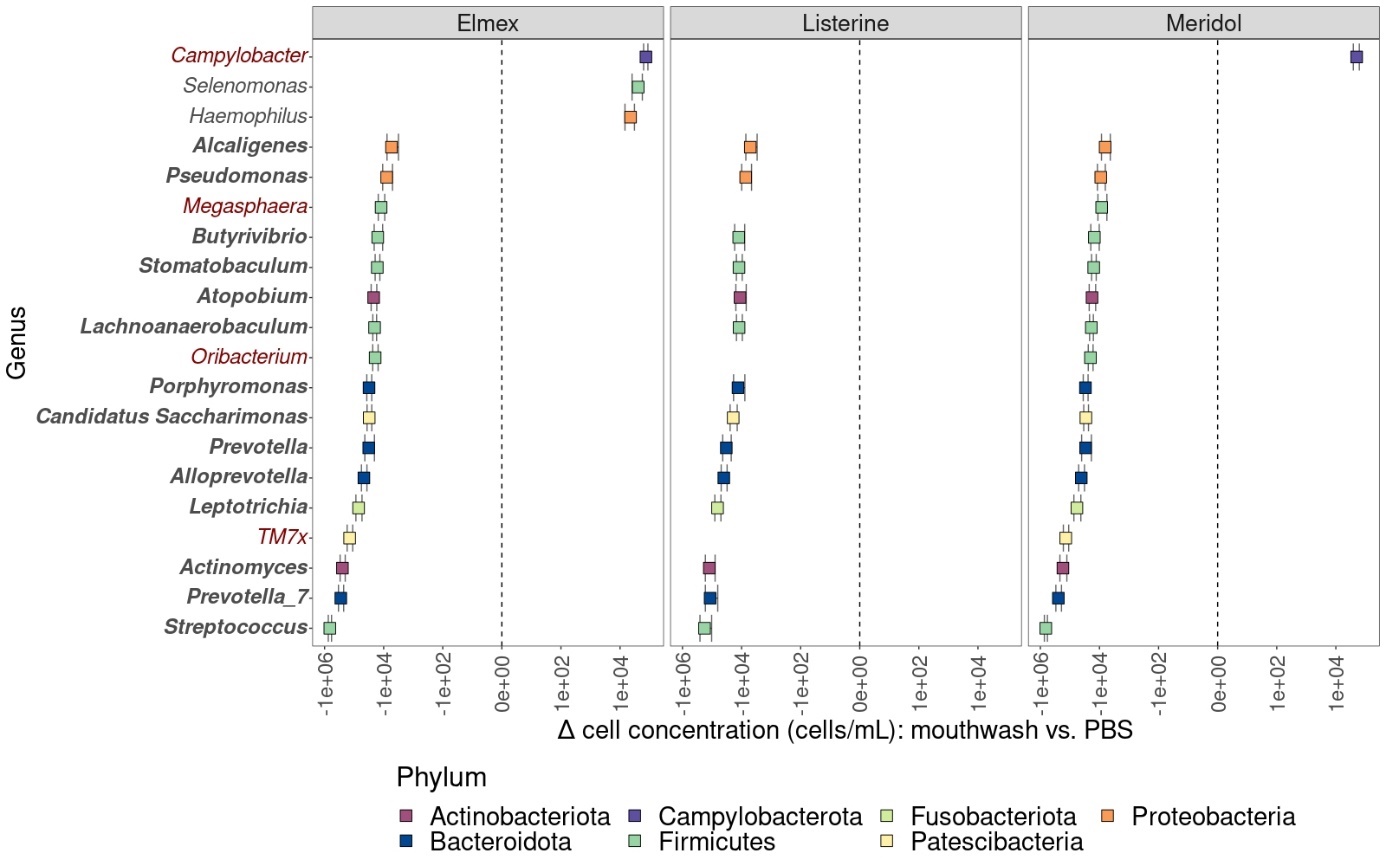

Figure S17. Differentially abundant genera for the different mouthwashes compared to the control (PBS) for the quantitative community data. Only genera with a relative abundance higher than 1% in at least one of the samples are shown in the figure. Genus names in bold indicate genera that were differentially abundant for all three mouthwashes and genus names in red indicate genera that were differentially abundant genera for both Elmex and Meridol. Error bars show the standard error.

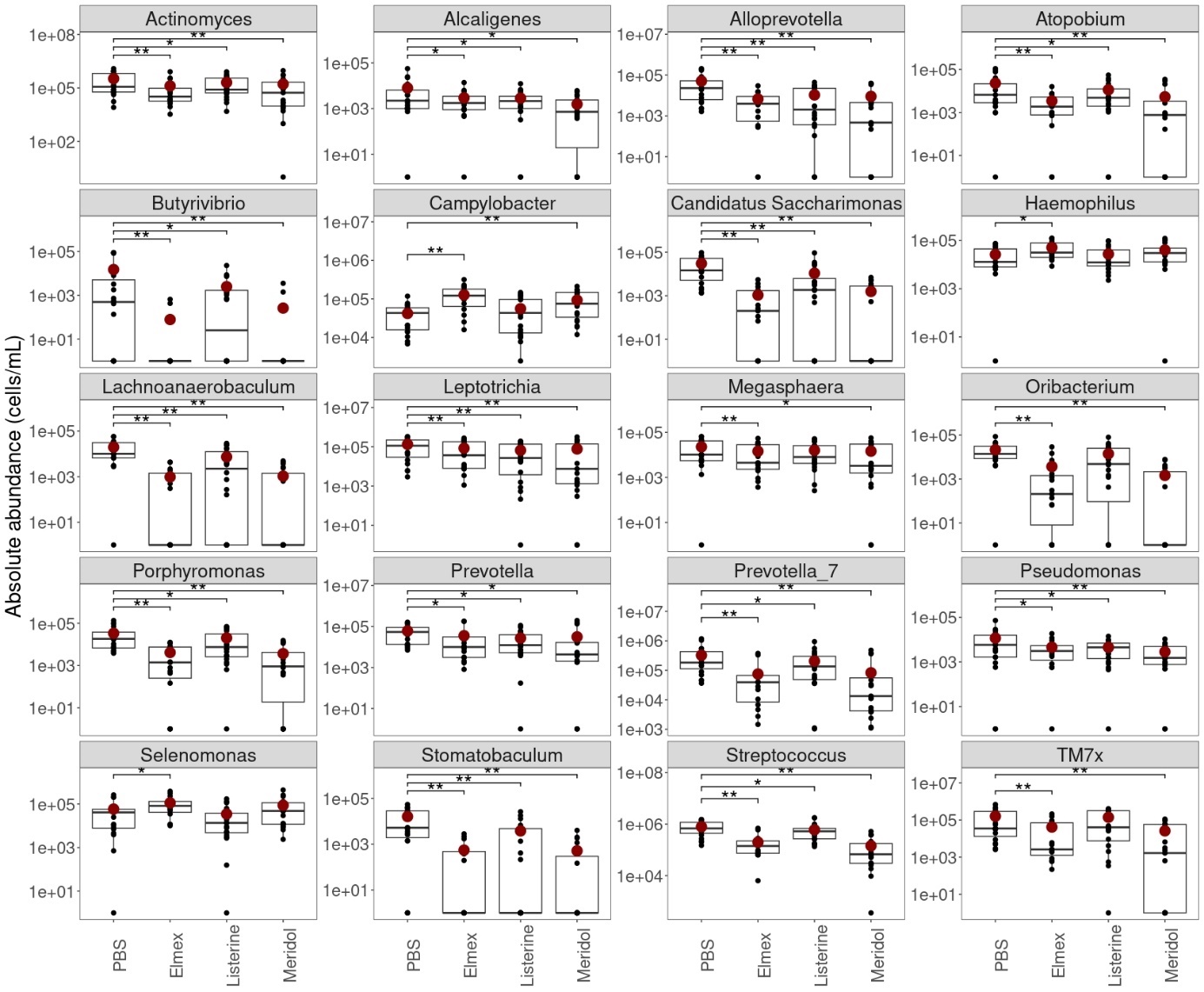

Figure S18. Absolute abundances of genera that were significantly different from the control (PBS) for at least one of the mouthwashes. Only genera with a relative abundance higher than 1% in at least one of the samples are shown. Red dots show the mean absolute abundance of the genus. Statistical differences were assessed using mixed-effect models with Benjamini-Hochberg correction for multiple testing and are indicated with asterisks (*P* < 0.05: *; *P* < 0.01: **).

**Supplementary 15. Organic acid concentrations after incubation by donor**

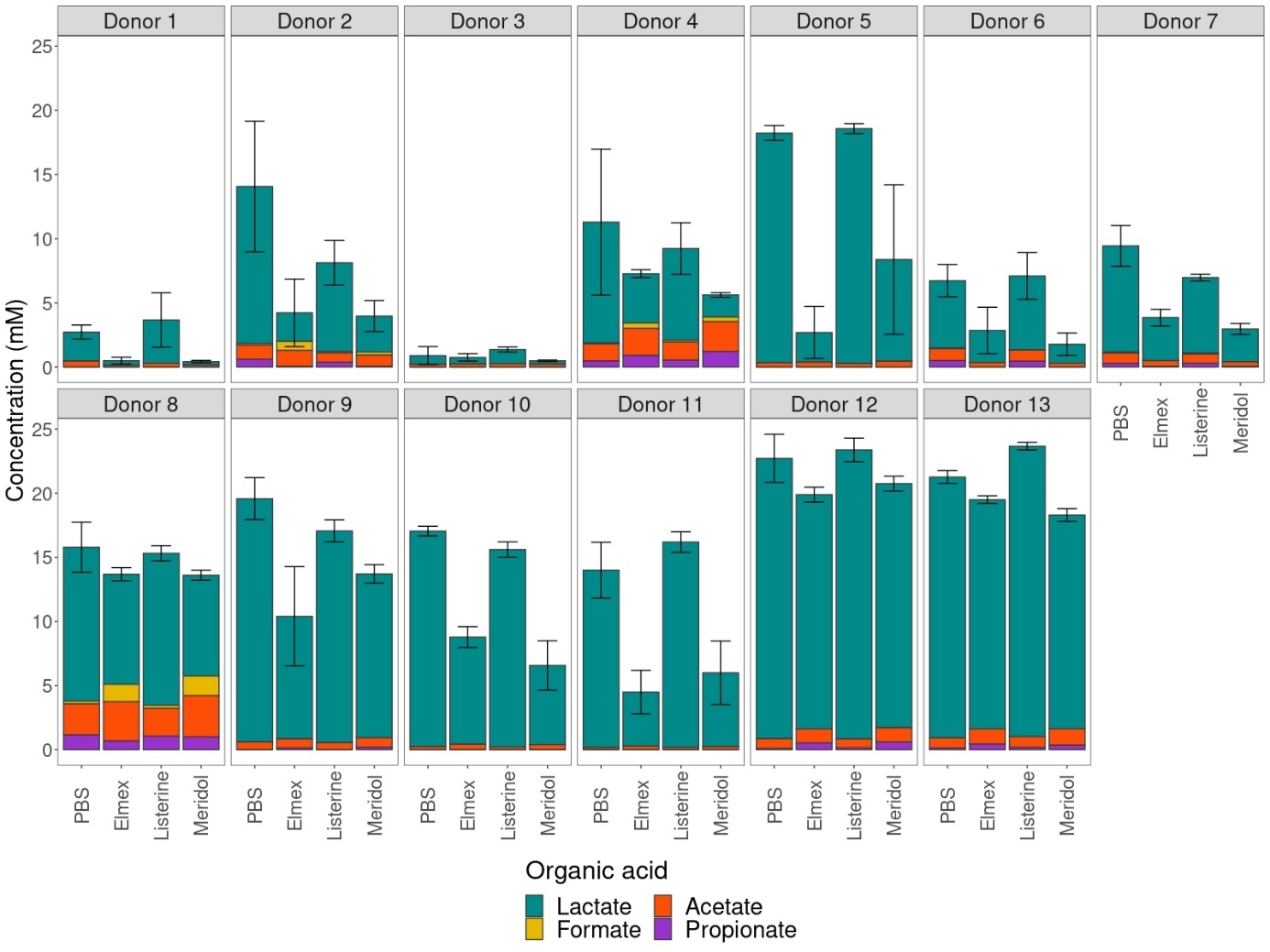

Figure S19. Molar organic acid concentrations after incubation of mouthwash-treated saliva in minimal medium with sucrose as carbon source for each donor. Error bars depict the standard deviation for the total organic acid concentrations.

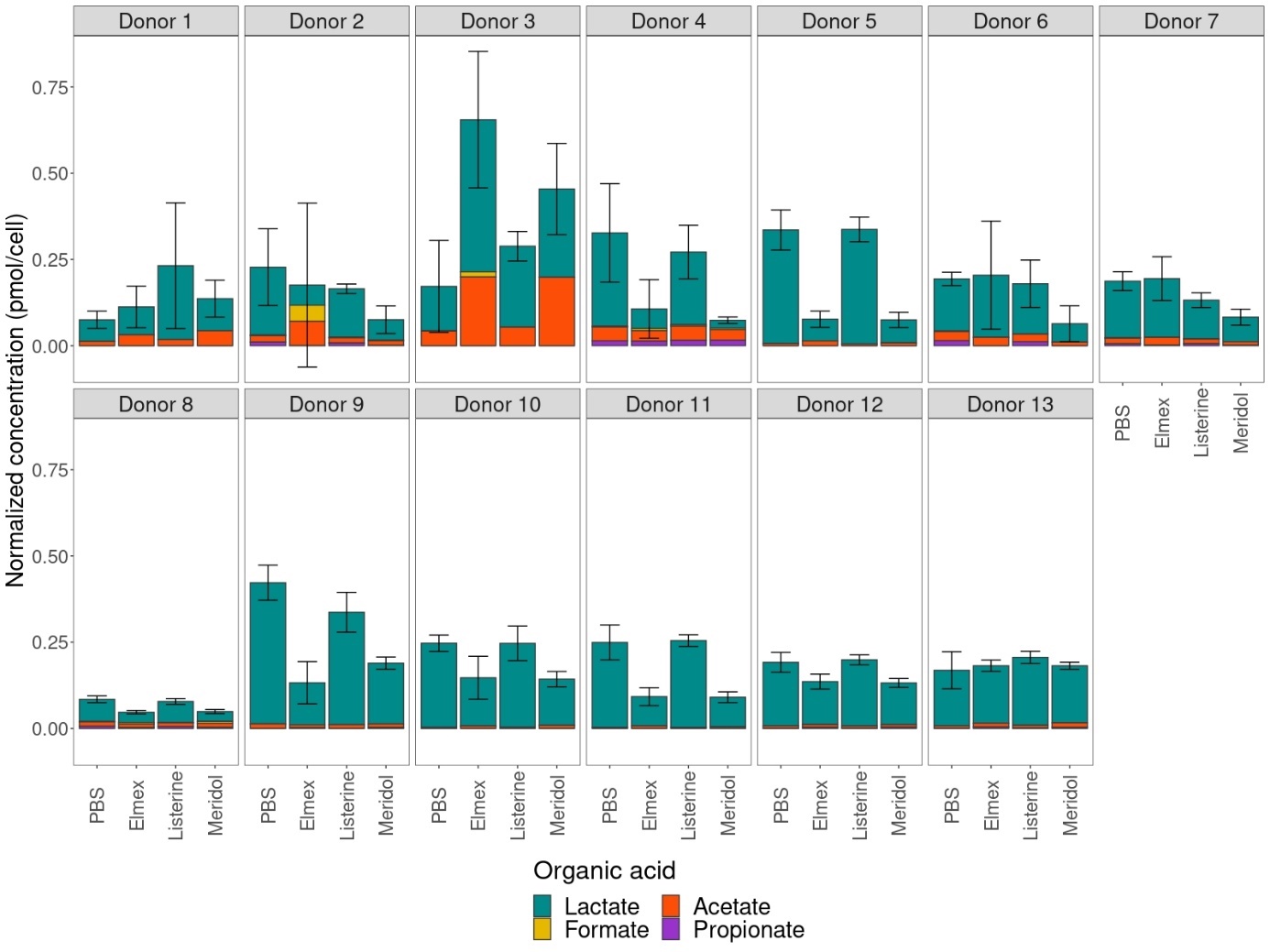

Figure S20. Total cell concentration normalized organic acid concentrations after incubation of mouthwash-treated saliva in minimal medium with sucrose as carbon source for each donor. The normalized concentrations were obtained by dividing the molar concentrations by the total cell concentration. Error bars show the standard deviation for the total normalized organic acid concentration.

**Supplementary 16. Community composition after incubation**

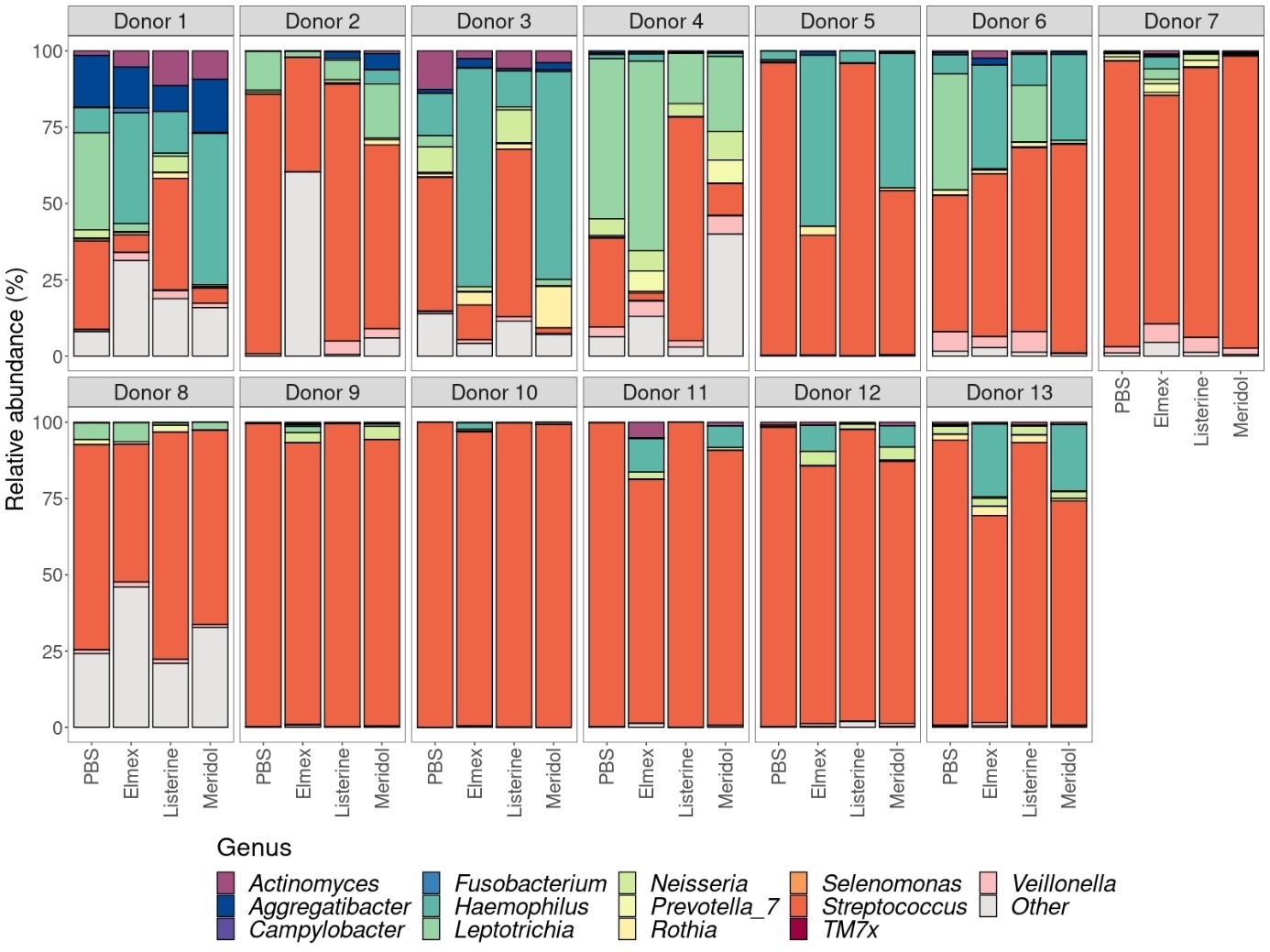

Figure S21. Relative abundance of the 13 most abundant genera after incubation. Lower abundant genera were grouped into ‘Other’.

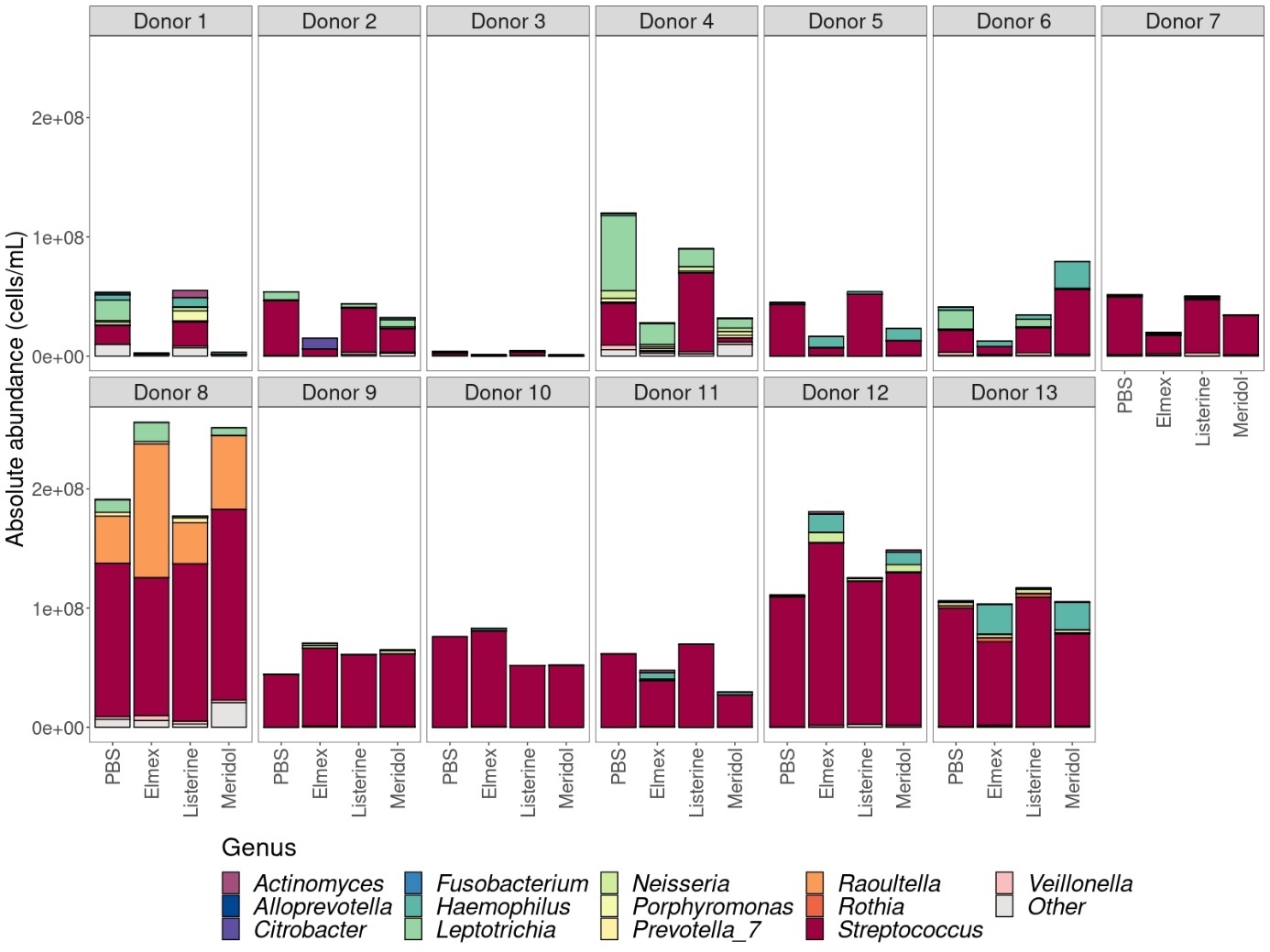

Figure S22. Absolute abundances of the top 13 most abundant genera after incubation. Lower abundant genera were grouped into ‘Other’.

**Supplementary 17. Differential abundance analysis after incubation**

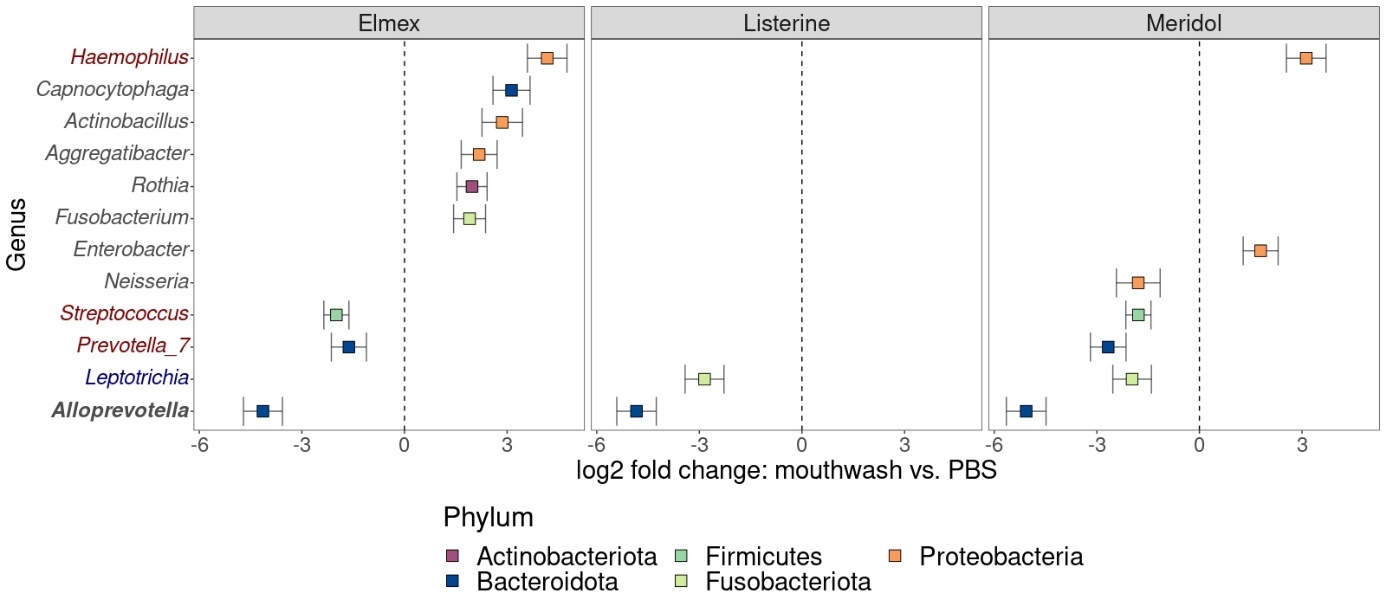

Figure S23. Differentially abundant genera compared to the control (PBS) for the proportional community composition of mouthwash treated saliva after incubation. Only genera with a relative abundance higher than 1% in at least one of the samples are shown in the figure. Genus names in bold indicate genera that were different in abundance in all three mouthwashes, genus names in blue indicate differentially abundant genera in both Listerine and Meridol, and genus names in red indicate genera that were differentially abundant for both Elmex and Meridol. Error bars show the standard error.

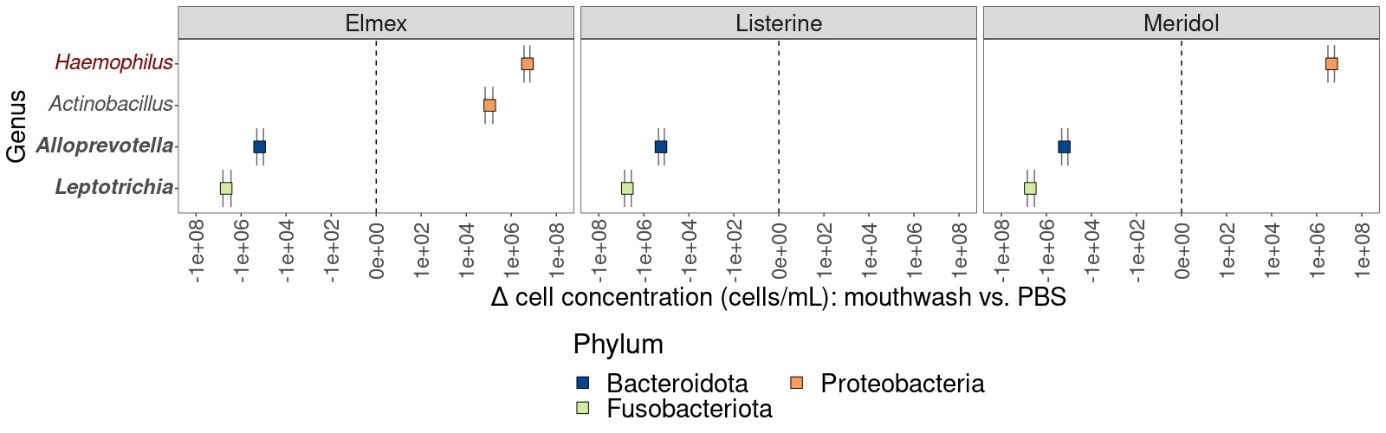

Figure S24. Differentially abundant genera compared to the control (PBS) for the quantitative community composition of mouthwash-treated saliva after incubation. Only genera with a relative abundance higher than 1% are represented in the figure. Genus names in bold indicate genera that were differentially abundant for all mouthwashes, and genus names in red indicate genera that were differentially abundant for both Elmex and Meridol. Error bars show the standard error.
